# Model Checking via Testing for Direct Effects in Mendelian Randomization and Transcriptome-wide Association Studies

**DOI:** 10.1101/2021.07.09.451811

**Authors:** Yangqing Deng, Wei Pan

## Abstract

It is of great interest and potential to discover causal relationships between pairs of exposures and outcomes using genetic variants as instrumental variables (IVs) to deal with hidden confounding in observational studies. Two most popular approaches are Mendelian randomization (MR), which usually use independent genetic variants/SNPs across the genome, and transcriptome-wide association studies (TWAS) using cis-SNPs local to a gene, as IVs. In spite of their many promising applications, both approaches face a major challenge: the validity of their causal conclusions depends on three critical assumptions on valid IVs, which however may not hold in practice. The most likely as well as challenging situation is due to the wide-spread horizontal pleiotropy, leading to two of three IV assumptions being violated and thus to biased statistical inference. More generally, we’d like to conduct a goodness-of-fit (GOF) test to check the model being used. Although some methods have been proposed as being robust to various degrees to the violation of some modeling assumptions, they often give different and even conflicting results due to their own modeling assumptions and possibly lower statistical efficiency, imposing difficulties to the practitioner in choosing and interpreting varying results across different methods. Hence, it would help to directly test whether any assumption is violated or not. In particular, there is a lack of such tests for TWAS. We propose a new and general GOF test, called TEDE (TEsting Direct Effects), applicable to both correlated and independent SNPs/IVs (as commonly used in TWAS and MR respectively). Through simulation studies and real data examples, we demonstrate high statistical power and advantages of our new method, while confirming the frequent violation of modeling (including IV) assumptions in practice and thus the importance of model checking by applying such a test in MR/TWAS analysis.

**Author Summary:** With the increasing availability of large-scale GWAS summary data of various complex traits/diseases and software packages, it has become convenient and popular to apply Mendelian randomization (MR) and transcriptome-wide association studies (TWAS), using genetic variants as instrumental variables (IVs), to address fundamental and significant questions by unraveling causal relationships between complex or molecular traits such as gene expression and other complex traits. However, the validity of such causal conclusions critically depends on the validity of the model being used, including three key IV assumptions. In particular, with the wide-spread horizontal pleiotropy of genetic variants, two of the three IV assumptions may be violated, leading to biased inference from MR and TWAS. This issue may become more severe as more trait-associated genetic variants are used as IVs to increase the power of MR and TWAS. Although there are some methods to check the modeling assumptions for MR with independent genetic variants as IVs, there is barely any powerful one for TWAS (or more generally for MR and similar methods) with correlated SNPs as IVs. We propose such a powerful method applicable to both MR and TWAS with local or genome-wide, possibly correlated, SNPs as IVs, demonstrating its higher statistical power than several commonly used methods, while confirming the frequent violation of modeling/IV assumptions in TWAS with our example GWAS data of schizophrenia, Alzheimer’s disease and blood lipids. An important conclusion is that in practice it is necessary to conduct model checking in MR and TWAS, and our proposed method is expected to be useful for such a task.

## 1 Introduction

It is of great interest in estimating and testing the causal effect of a risk factor/exposure X on an outcome Y. However, for observational data, due to the presence of unmeasured confounders, say U, it is difficult to tell whether an observed association between X and Y really indicates a causal relationship. Mendelian randomization (MR) has been applied as a popular and powerful approach to addressing this problem for causal inference, using genome-wide significant genetic variants, typically single-nucleotide polymorphisms (SNPs), as instrumental variables (IVs). Various versions of MR have been proposed, most of which are convenient to implement since they only require the use of GWAS summary data. Recently, MR has been widely applied to obtain substantial findings, and one example is Richardson et al. (2019), which found significant evidence for the causal relationships between many traits of interest by utilizing the large-scale UK Biobank data (Sudlow et al. 2015; Neale Lab 2017). A commonly used workflow of MR (or TWAS) analysis is illustrated in Figure S1 in the supplementary materials. However, as expected, the validity of any MR analysis critically depends on its modeling assumptions, in particular including three modeling assumptions on valid IVs; an IV has to satisfy the following three conditions to be valid, as depicted in Figure 1 (A):

1. The IV is associated with the exposure X.
2. The IV, conditional on X, is not directly associated with the outcome Y.
3. The IV is not associated with the hidden confounder U.

**Figure 1.**
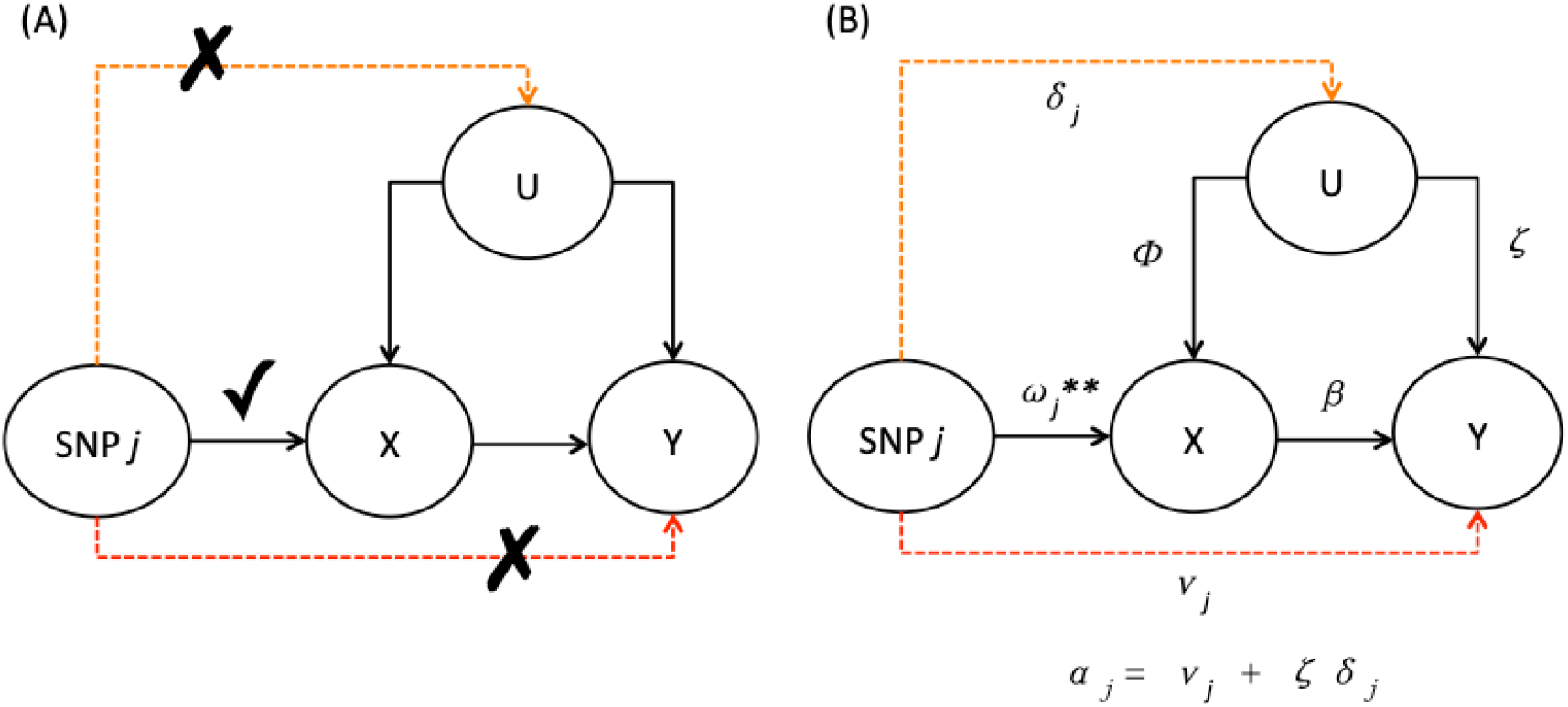
Illustrative diagrams for (A) valid IV assumptions and (B) horizontal pleiotropic effects violating two of the three IV assumptions.

When any of the three assumptions is violated, the conclusion from MR can be incorrect. Among the three assumptions, the first one appears easiest to handle: one can simply ensure that a SNP/IV is indeed associated with the exposure X by using a stringent (genome-wide) significance level; the challenge lies in the other two, especially the third one with unobserved confounding. It is well known that, the wide-spread horizontal pleiotropy (Cotsapas et al. 2011; Wang et al. 2015; Verbanck et al. 2018), i.e. when an SNP is associated with multiple traits (e.g. X and Y here) through different pathways, will likely result in the violation of one or both of the last two IV assumptions, leading to invalid IVs and thus incorrect causal conclusions. For instance, horizontal pleiotropy can occur in the form of having a pathway from the IV directly to Y, which corresponds to **uncorrelated** pleiotropy violating the second IV assumption. It can also happen when there is an additional pathway from the IV to U then to Y, corresponding to **correlated** pleiotropy (Morrison et al. 2020) and violating the third IV assumption. In this paper, we use horizontal pleiotropy to refer to both cases; see Fig 1(B). We also note that violation of the third IV assumption may not always be due to horizontal (correlated) pleiotropy when, for example, the confounding is due to population stratification (i.e. when the effect direction is from U to IVs, instead of from IVs to U); As shown in the Supplementary, this may also imply some “direct effects” of the IV on Y, not mediated through X, for which our proposed methods appear to be applicable, though this is beyond the scope of this paper and needs further investigation.

In order to reduce the negative impact of invalid IVs, some MR methods have been developed to account for horizontal pleiotropy (with GWAS summary data) (Bowden et al. 2015; Bowden et al. 2016; Burgess et al. 2016a; Burgess et al. 2016b; Kang et al. 2016; Windmeijer et al. 2016; Burgess et al. 2018; Hartwig et al. 2017; Verbanck et al. 2018; Zhao et al. 2018; Burgess et al. 2019; Jiang et al. 2019; Qi & Chatterjee 2020; Morrison et al. 2020; Xue et al. 2021). However, simulation studies have shown that it is unlikely that any method can completely solve the problem in all scenarios while possibly imposing its own other modeling assumptions (Slob et al. 2020; Xue et al. 2021). For example, the constrained instrumental variable (CIV) methods by Jiang et al. (2019) use a framework where each genotype’s effect on the outcome has to either go through the exposure or a pleiotropic phenotype that is observed, while in reality there can be many other unknown pathways. In addition, due to their own modeling assumptions and often much lower estimation efficiency, they may give quite different results, casting doubt on which results are valid.

Hence, it is important to detect whether any modeling or IV assumptions are violated before accepting MR results that may be problematic. For MR, one can apply Cochran’s Q or Rucker’s Q’ statistic for model checking, and MR-Egger to test for directional pleiotropy using its intercept term (Bowden et al. 2018). Dai et al. (2018) proposed a new method called global and individual tests for direct effects (GLIDE), which has been shown to have higher power than MR-Egger, but it has not been demonstrated to outperform Cochran’s Q or Rucker’s Q’ statistic (Bowden et al. 2018). Also, GLIDE seems to require individual level GWAS data for the outcome, which are often unavailable. Most importantly, all of these methods can only be applied to independent IVs, excluding their use with correlated IVs as in transcriptome-wide association studies (TWAS).

TWAS has been proposed recently to examine the relationship between a gene’s expression and an outcome (Gamazon et al. 2015; Gusev et al. 2016). As in MR, if the three IV assumptions hold, such a detected association implies a causal relationship. TWAS is a two-stage least squares (2SLS) regression approach in the framework of IV regression; some correlated cis-SNPs near a gene are used as IVs to impute or predict the gene’s expression level. As MR, an advantage of TWAS is that it can be conducted with GWAS summary data. Here we refer to TWAS in a general sense as an MR extension that may include any other trait as an exposure while using correlated (cis- or whole genome-wide) SNPs as IVs. Some authors have found that TWAS can gain power over MR by more effectively using correlated SNPs, instead of independent ones, as IVs (Knutson and Pan 2020). Nevertheless, cautions have to be taken when interpreting TWAS results because TWAS, as MR, may suffer from using invalid IVs (Mancuso et al. 2019; Wainberg et al. 2019; Wu and Pan 2020). The standard/default TWAS only models the relationship between an outcome trait and a gene’s predicted expression, which means that possible horizontal pleiotropy is not considered and thus can impact the result. To handle pleiotropy, Barfield et al. (2017) proposed a method called LD-aware (LDA) MR-Egger, which models direct/pleiotropic effects as random effects as in MR-Egger, but differs from the latter in modeling the joint effects, not marginal effects, of the SNPs/IVs. This approach can handle certain situations much better than the standard TWAS, but it may still have problems with invalid IVs, especially when the InSIDE (Instrumental Strength Independent of Direct Effect) assumption is violated; in the presence of correlated pleiotropy violating the third IV assumption, the InSIDE assumption will be violated. This means even when relatively robust methods like MR-Egger are used, it is still important to test for pleiotropic effects, or more generally, for the goodness-of-fit of the model being used, which will influence the validity of the results. Like MR-Egger, LDA MR-Egger itself can be used to test whether there is directional (uncorrelated) pleiotropy by testing the intercept term, but it cannot handle correlated pleiotropy while its power is quite low as to be demonstrated. Hence, to ensure valid conclusions from TWAS, model checking, including testing for the presence of (uncorrelated and/or correlated) pleiotropic effects for the violation of the second and third valid IV assumptions, is much needed in TWAS.

Several methods, including colocalization tests, have been proposed to test for pleiotropy, but they may not be able to distinguish horizontal pleiotropy from vertical pleiotropy. For instance, SMR (with the HEIDI test) of Zhu et al. (2015) can integrate summary level GWAS and eQTL data to find genetic variants with pleiotropic effects on both the GWAS trait and gene expression, but it cannot distinguish whether a variant is affecting the GWAS trait and gene expression through two different pathways (i.e., horizontal pleiotropy), or instead, it effects the GWAS trait through the gene as a mediator (i.e., vertical pleiotropy). Since only horizontal pleiotropy is problematic to the IV assumptions, it is important to test on horizontal pleiotropy directly, which is the goal here.

In consideration of the above limitations and challenges, there is a strong need for a LDA (LD-aware) method that can test for horizontal pleiotropy with higher power using possibly correlated IVs, which can be either some local or genome-wide SNPs. Yuan et al. (2020) proposed a likelihood-based method that aims to test and control for horizontal pleiotropy. Their models are more general than the LDA MR-Egger approach, but their method applies the burden test on the perhaps over-simplified assumption that the horizontal pleiotropic effects of the IVs/SNPs are all equal, which can be violated, leading to power loss. To address the above issues, we propose a general GOF test, called TEDE (TEsting Direct Effects), for model checking in MR and TWAS; it can detect violations of modeling assumptions, including IV assumptions, though testing for the presence of direct effects of the IVs on the outcome, which, as depicted in Figure 1(B) and Supplementary Figure S2(B), can be due to the violation of the second or third IV assumption, and population structure, or to other reasons to be discussed later. We propose two versions, TEDE-Sc (TEDE-Score) by applying the score test, and TEDE-aSPU by an adaptive test called the aSPU test that is particularly powerful for a large number of SNPs/IVs (Pan et al. 2014). The new tests can be seamlessly applied to both MR and TWAS. We also propose a way to control type I error rates better by taking into consideration of the variance of an estimated SNP-exposure association. Through simulation studies we show that our new methods are able to handle both uncorrelated and correlated IVs (for MR and TWAS respectively), and more importantly, have higher power than Cochran’s Q-statistic (for MR) and LDA TWAS-Egger while satisfactorily controlling type I errors. We apply the methods to a large GWAS dataset of schizophrenia (SCZ) (Schizophrenia Working Group of the Psychiatric Genomics Consortium 2014) and the GWAS data of various traits based on the imputed UK Biobank data (Sudlow et al. 2015; Neale Lab 2017) to further demonstrate how different methods perform in the context of MR. We also apply the methods to the ADNI data (Shen et al. 2014), the IGAP stage 1 AD data (Lambert et al. 2013) and the lipid data (Teslovich et al. 2010; Willer et al. 2013) to show the advantages of our new methods in TWAS, while confirming the commonality of the violation of modeling/IV assumptions perhaps due to the wide-spread horizontal pleiotropy in reality.

## 2 Methods

### 2.1 Existing Model Checking Methods in MR

In this section, we give a brief review of some representative GOF tests in MR with GWAS summary data. We only include the most popular Cochran’s Q test and MR-Egger, because Rucker’s Q’ test performs similarly to Cochran’s Q while GLIDE requires individual-level data for the outcome (Bowden et al. 2018; Dai et al. 2018), which is often unavailable in practice. For simplicity, we only consider linear regression models throughout this paper. Although logistic regression models are often considered for binary outcomes, their use in instrumental variable regression for causal inference is challenging and complicated; on the other hand, due to small effects of SNPs/IVs, linear models can approximate logistic models well (Xue et al. 2020) and thus have been widely used in MR and TWAS.

Suppose we have *p* independent SNPs. 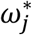 is the total effect of SNP *j* on X, while 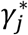 is the total effect of SNP *j* on Y based on marginal models (i.e. X ~ SNP *j* ~ SNP *j*). Denote 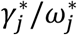 by 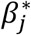, the real X to Y effect by *β* and the SNP to Y direct effect by *α_j_*. For instrumental variable analysis, as shown in Figure 1 (A), valid IV assumptions require that there is no direct effect from the IV (SNP *j*) to the confounders or to the outcome. If there is an effect from SNP *j* to the outcome not mediated through X, we say there is **horizontal pleiotropy**, and its presence leads to the violation of the last two IV assumptions and thus biased MR analysis. Figure 1 (B) shows the notations of different effects in the possible presence of horizontal pleiotropy. Here *α_j_* is the total SNP to Y effect that does not go through X, which consists of two parts: the direct effect *v_j_* of the SNP on Y that does not go through U, and the effect *ζδ_j_* from the path going through U. In this sense, we can combine the violation of valid IV assumptions (2) and (3) into one condition, *α_j_* ≠ 0, and we only need to test whether *α_j_*=0. Based on Figure 1 (B), we know that 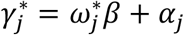, and thus 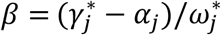; note that, we use 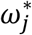 here to represent 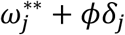 in Figure 1(B). No horizontal pleiotropy means that each *α_j_* is 0 and each 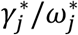 is equal to *β*, and thus we can test 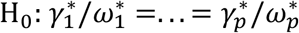 or 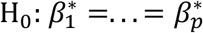, which only needs the marginal summary statistics 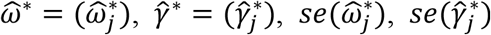.

It is noted that the problem we consider is more general than horizontal pleiotropy (that is the focus here): the presence of the direct effects *α_j_* ≠ 0 can arise due to the violation of other modeling assumptions. For example, as shown in Supplementary Figure S2(B), population stratification can also cause the violation of the IV assumptions manifested as the presence of some direct effects *α_j_*.

#### 2.1.1 Cochran’s Q Test

As a general GOF test, Cochran’s Q test uses the test statistic

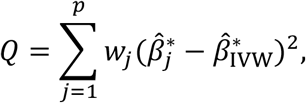

where 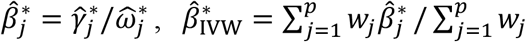 and 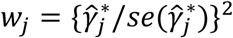. If the null hypothesis is true (i.e. if the model fits the data well, including that there is no global pleiotropy), *Q* should follow 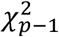 (Bowden et al. 2018).

#### 2.1.2 MR-Egger

Another way to test horizontal pleiotropy is to apply the MR-Egger method (Bowden et al. 2015) and examine the intercept term. The model is

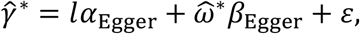

where *ε~N*(0, Σ) and *l* = (1…1)′, *α*_Egger_ and *β*_Egger_ are two parameters. Here Σ is usually assumed to be a diagonal matrix with diagonal elements being 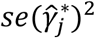’s. To test pleiotropy, we simply test whether *α*_Egger_ = 0. Note that *α*_Egger_ models the average direct effect of the SNP to Y, and thus testing *α*_Egger_ = 0 is actually testing whether there is *directional* pleiotropy (i.e. the mean of the direct effects is nonzero). Also note that for MR-Egger, the alleles of the variants are flipped before fitting the model to ensure that 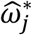’s are all positive.

### 2.2. Model Checking in TWAS

In this section, we introduce some LDA methods for model checking, including testing for horizontal pleiotropy, applicable to both TWAS and MR. TWAS examines the effect of a gene’s expression (X) on the outcome (Y) using the gene’s cis-SNPs as IVs in two stages. In the first stage, X is regressed on and thus predicted by the SNPs. In the second stage, Y is regressed on the predicted X with the regression coefficient as the key parameter of interest, measuring the effect of X on Y. This procedure works well when the modeling assumption hold, including that valid IV assumptions are not violated. However, if there is horizontal pleiotropy or population structure (as depicted in Figure 1(B) and Supplementary Figure 2(B)), some of the SNPs may have direct effects on Y that are not mediated through X, under which the above standard TWAS (i.e. only regressing Y on the predicted X) may lead to biased inference on the causal relationship between X and Y. Hence, it is equally important to check the TWAS model via testing for direct effects of the IVs.

Note that, even though both TWAS and MR appear similar as two-stage least squares IV regression in inferring the causal effect of X on Y using some SNPs as IVs, typical MR methods use independent SNPs as IVs while TWAS uses a gene’s local/cis SNPs that are usually correlated (i.e. in linkage disequilibrium, LD), explaining why LDA methods are needed for the latter. Here we use TWAS to also refer MR extensions with correlated SNPs (either locally or across a whole genome). In TWAS we model the joint effects *ω_j_* (based on the joint model X ~ multiple SNPs), instead of the marginal effects 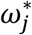 (on the marginal models X ~ one SNP). This is crucial because *ω_j_* is usually quite different from 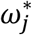 when the SNPs are correlated, and *ω_j_* captures the effect of each SNP on X after adjusting for other SNPs.

Using notations similar to the previous section, suppose we have *p* SNPs, which can be correlated in our new settings, and their direct effects on Y are denoted by *α_j_*’s. *ω_j_* is the effect of SNP *j* on X, *γ_j_* is that of SNP *j* on Y based on joint models (i.e. X ~ SNPs, Y ~ SNPs). We can test 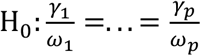. In practice, we are often provided with only GWAS marginal summary statistics 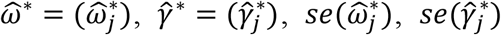. Using these with a reference panel, we can get estimates 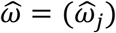 and 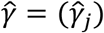, their standard errors 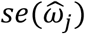 and 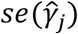, as well as 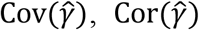 and 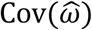 (Yang et al. 2012; Deng and Pan 2017).

#### 2.2.1 LDA MR-Egger

Barfield et al. (2018) proposed a method called LDA MR-Egger that applies the idea of Egger regression to TWAS while accounting for the LD structure of the SNPs. The model is

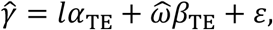

where *ε*~*N*(0, Σ) and *l* = (1…1)’, *α*_TE_ and *β*_TE_ are two parameters. Usually Σ is simply estimated as 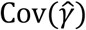. For our current problem, since we are interested in testing H_0_: *γ*_1_/*ω*_1_ =…= *γ_p_/ω_p_*, we can test *α*_TE_ = 0. Transform the model to make the error term follow *N*(0,1):

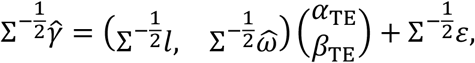

where 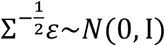. As a result, we can estimate *α* and *β* by

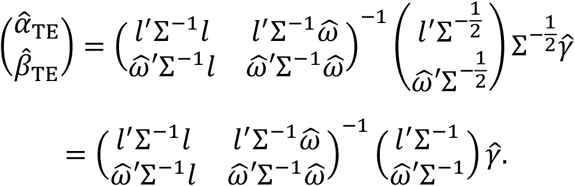

This is the same as the LDA MR-Egger estimate in Barfield et al. (2018). The covariance matrix is

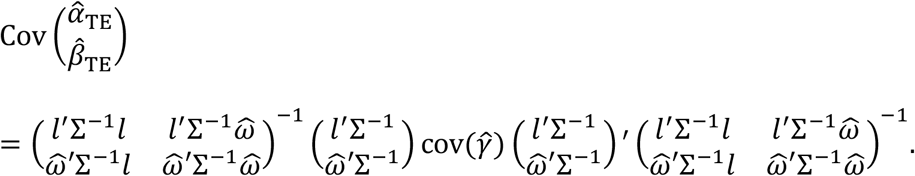

If we assume 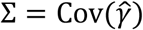, then

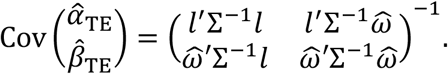

Thus we can get 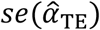 and 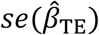. We use 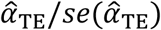 to test horizontal pleiotropy and 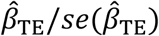 to test the effect of X on Y. Note that to be consistent with MR-Egger, we flip the alleles to make sure 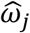’s are positive before estimating 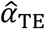 and 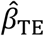.

#### 2.2.2 New Method: Its Framework

Our new method was motivated as a general GOF test for the original/standard TWAS. Suppose *X_i_* and *Y_i_* are the exposure and outcome for subject *i*, *G_i,j_* is the *j*th SNP/IV for subject *i*; all have been centered at 0. In Stage 1 of TWAS, we fit a linear model

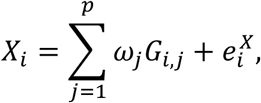

yielding estimates 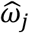’s and imputed/predicted gene expression/exposure 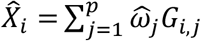. Then in Stage 2 we fit

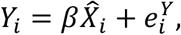

and thus estimating the causal effect *β* of X on Y, the parameter of interest in TWAS. Note that we do not include an intercept term in either model since the variables have already been centered at 0. This procedure only looks at the part of X and Y that is explained by the SNPs, which is able to account for hidden confounding when valid IV assumptions are met, and thus gives a good estimate of *β* in the presence of hidden confounding (i.e. U is present but not explicitly included in the models). However, as depicted in Figure 1(B) and Supplementary Figure S2(B), due to the violation of the second or third IV assumption, or to population structure, there will be non-zero direct effects *α_j_*’s on Y, leading to a mis-specified model being used in TWAS Stage 2. Accordingly, we propose a general GOF test for TWAS by testing whether the direct effects *α_j_*’s are all 0 or not in an expanded model

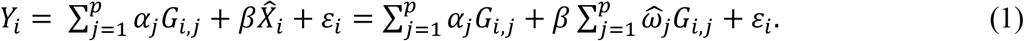

Then we test H_0_: *α*_1_ =…= *α_p_* = 0. If *H*_0_ is rejected, then the TWAS Stage 2 model is incorrect (in assuming no direct effects), possibly due to the violation of some modeling assumptions, including the presence of invalid IVs (e.g. due to horizontal pleiotropy) or of population structure. Furthermore, other violations of modeling assumptions can lead to rejecting H_0_, e.g. due to an incorrect model or bad estimation in Stage 1: if 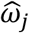 and *ω_j_* are quite different, even if other TWAS modeling/IV assumptions hold, it may still lead to a nonzero “direct effect” *α_j_*, and thus the rejection of H_0_.

It is noted that Model (1) is over-specified for parameter estimation, but testing on H_0_ is possible, including parameter estimation under H_0_. Our proposed testing framework is related to the Sargan test for over-identifying restrictions in IV regression (Sargan 1958; Windmeijer et al. 2019), and can be regarded as an extension to (2-sample) TWAS with GWAS summary data.

#### 2.2.3 New Method: TEDE

To test H_0_: *α*_1_ =…= *α_p_* = 0, we can use the score test or another test. We assume that *ε_i_*’s follow an i.i.d normal with mean 0 and variance 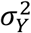. Note that this model is different from the true model since it uses 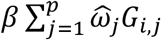, rather than 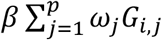, which may have potential problems if 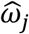’s are inaccurate and *β* is nonzero. We will demonstrate this further in our simulation studies. Denote the parameter vector by **θ** = (*α*_1_,…, *α_p_,β*)′ and the score vector by **U**(**θ**) = (*U*_1_(**θ**),…, *U*_*p*+1_(**θ**)). The log-likelihood is

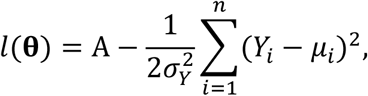

where 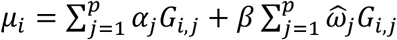 and A is a constant that does not involve **θ**.

For *j* = 1,…, *p*, we have

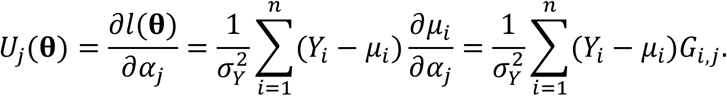

We also have

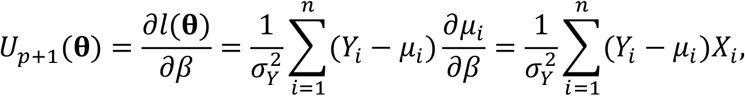

where 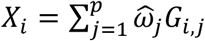. To apply the score test, we need to estimate 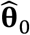, which is the MLE of **θ** under H_0_: *α*_1_ =…= *α_p_* = 0. With *α*_1_ =…= *α_p_* = 0, we know 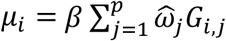 and

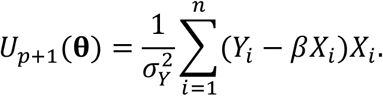

By setting *U*_*p*+1_(**θ**) = 0, we get 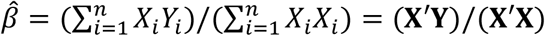, where **X** = (*X*_1_,…, *X_n_*)′. It is easy to see that 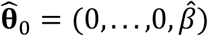 maximizes *l*(**θ**) under H_0_.

As a result, we know 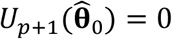 and

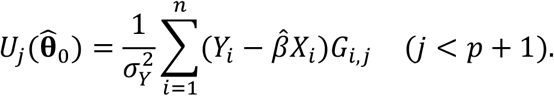

To estimate the covariance matrix of 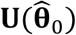, we need to calculate

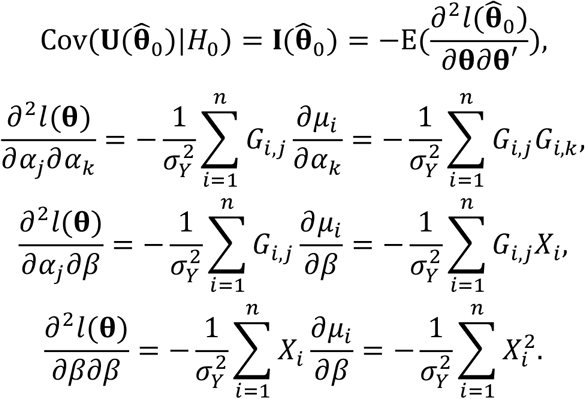

Hence, by denoting **G** = (**G**_1_,…, **G**_*p*_), **G**_*j*_ = (*G*_1,*j*_,…, *G_n,j_*)′, we can obtain

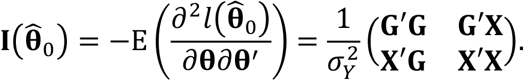

##### TEDE-Sc

We can test H_0_ since we know that 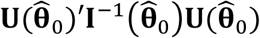 asymptotically follows a chi-squared distribution with degrees of freedom equal to the rank of 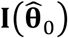 under H_0_. We call this test TEDE-Sc (TEsting Direct Effects by the Score test). Note that its test statistic requires an estimate of 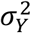. Under H_0_, 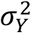 can be easily estimated as the sample variance of 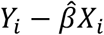, which means 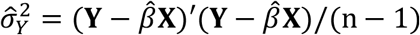. We can also calculate 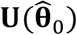 and 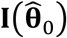 with GWAS summary statistics and a genotypic reference panel, since they allow us to estimate 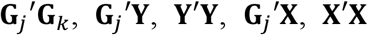 and **X′Y** (Deng and Pan 2017).

##### TEDE-aSPU

Denote 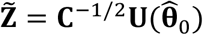, where **C** is a diagonal matrix with the same diagonal elements as 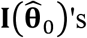. 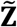 can be regarded as U-scores standardized by their standard errors. Under H_0_, 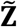 should asymptotically follow 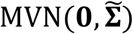 where 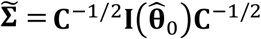, and thus 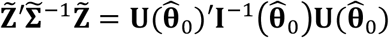 can be used to test H_0_, which is exactly the same as TEDE-Sc. To test whether all of the scores in 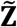 are 0, which tells whether H_0_ is true, we can apply the SPU tests and aSPU test (Pan et al. 2014). First, we denote 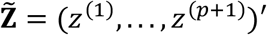 and define

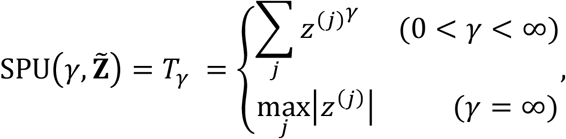

where *γ* is usually chosen from {1, 2,…, 8, ∞}. We sample 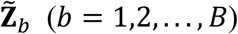 from the null distribution 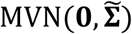, and the p-value for the SPU test is

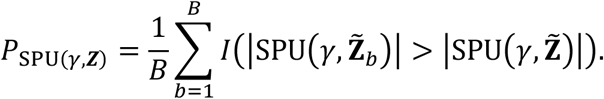

The general idea is to simply look at whether the sum of powered scores is too extreme, since under H_0_, the scores should have mean 0 and their powered sum should not be too large. If we look at a set of different *γ*’s, denoted by Γ = {*γ*_1_, *γ*_2_ … *γ_r_*}, each of them yields a different p-value *P*_SPU(*γ_t_*,**Z**)_. To combine these results, we define the aSPU test statistic as 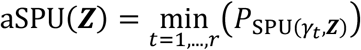. For each power index *γ_t_*, we calculate

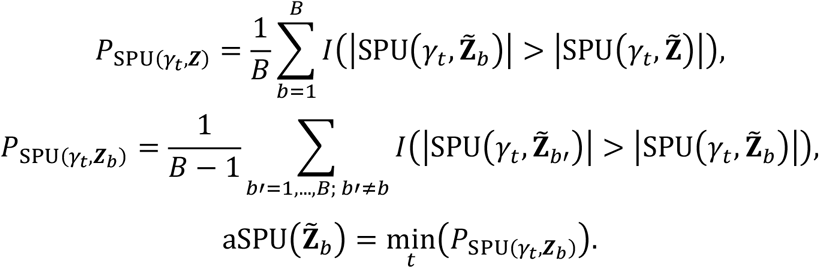

The p-value of the aSPU test is calculated as 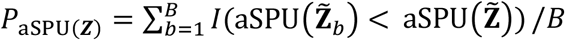. We call this approach TEDE-aSPU, which applies the aSPU test to our problem of testing invalid IVs,

The SPU(1) and SPU(2) tests, corresponding to the burden test and a variance component/kernel test respectively, have been shown to have higher power when the signals are dense (i.e. more *α_j_*’s are nonzero), whereas SPU(8) and SPU(∞) usually works better when the signals are sparse (i.e. few *α_j_*’s are nonzero) (Pan et al. 2014). The aSPU test is able to combine their strengths and thus perform well in various scenarios. Since TEDE-Sc and SPU(2) both look at the second order of the scores (one with 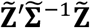, the other with 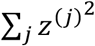), we expect TEDE-Sc to also perform better when the invalid IVs are relatively dense. When the invalid IVs are sparse or high-dimensional, we expect that TEDE-aSPU to be more powerful.

Our method does not require the InSIDE (Instrument Strength Independent of Direct Effect) assumption to hold; we will demonstrate through simulations that our method performed well even when InSIDE was violated. It is also worth noting that our previous model and results are based on the assumption that 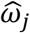’s are fixed values. In reality, what follows i.i.d. normal with mean 0 and 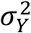 under the null should be 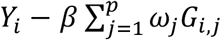 instead of 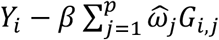. As a result, by ignoring the estimation error and variability of 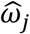’s, we may inaccurately estimate *β* and *α_j_*’s, which may lead to inflated type I errors (if, as default in this paper, one does not really care about the estimation errors of *ω_j_*’s but only the functional form of the specified model; otherwise it would be power, instead of type I error). This may happen for TWAS since usually the sample size in the first stage is not large enough to ensure the accuracy of 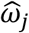’s. To mitigate this issue, we can incorporate the variance of 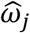’s by replacing 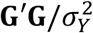 with 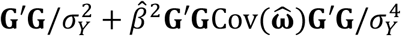 when calculating 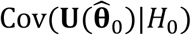, since the first *p* elements of 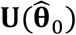 under *H*_0_ can be written as 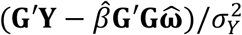. We do not need to worry about the other elements in **S** because 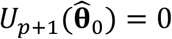 and those elements will not have any effect on TEDE-Sc or TEDE-aSPU. We call the approach with the modified covariance estimate TEDE-aSPU2, which is expected to better control type I errors. We can also use the modified covariance estimate in TEDE-Sc, and we call this approach TEDE-Sc2. Also note that when the significance threshold is too small (e.g. <5e-8), using the original version of aSPU with summary statistics may take too much time. Alternatively, we can apply the aSPU test based on either its asymptotics (Xu et al. 2016) or on importance sampling (Deng et al 2020), which has been shown to perform well when p is large and small respectively.

#### 2.2.4 Connection between TEDE-aSPU and TWAS-aSPU

A more powerful association test in TWAS has been proposed by Xu et al. (2017). Their model is

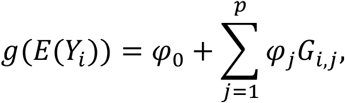

where the link function *g*() is the identity function for quantitative trait *Y_i_*. For the linear case with variables already centered at 0, the model becomes 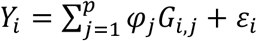.

To assess possible association between the trait and the SNPs, one applies the aSPU test to the null hypothesis *H*_0_: *φ*_1_ =…= *φ_p_* = 0. Comparing this model to our working model (1), we can decompose each 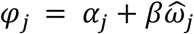. It is clear why this test was shown to be more powerful than TWAS: it tests not only on the causal effect *β* of X on Y (as does TWAS), but also on direct effects of the SNPs. In other words, the association test consists of two components: one is for causal effects of X on Y as in TWAS, and another on the mis-specified TWAS model (e.g. due to invalid IVs) as aimed by TEDE proposed here.

#### 2.2.5 Applying LDA Methods to MR

For MR analysis with independent SNPs, we can directly apply the LDA methods, including TEDE, for model checking or GOF testing because eventually they are all testing H_0_: *α*_1_ =…= *α_p_* = 0. As long as we know the MAF of each SNP (either from a GWAS summary dataset or a reference panel), we can calculate the variance for each SNP, leading to a diagonal LD covariance matrix. If MAF information is already provided in the GWAS summary data, we do not need to use a reference panel.

### 2.3 Data Availability

The ADNI data were obtained from the Alzheimer’s Disease Neuroimaging Initiative (ADNI) database (adni.loni.usc.edu). For up-to-date information, see www.adni-info.org. The SCZ data (Schizophrenia Working Group of the Psychiatric Genomics Consortium 2014) is obtained from https://www.med.unc.edu/pgc/results-and-downloads. The IGAP data (Lambert et al. 2013) is publicly available at http://web.pasteur-lille.fr/en/recherche/u744/igap/igap_download.php. The 2010 (Teslovich et al. 2010) and 2013 lipid data (Willer et al. 2013) are also publicly available at http://csg.sph.umich.edu/abecasis/public/lipids2010 and http://csg.sph.umich.edu/abecasis/public/lipids2013/, respectively. The GWAS results using imputed UK Biobank data (Sudlow et al. 2015; Neale Lab 2017) are provided at http://biobank.ctsu.ox.ac.uk/crystal/. The FUSION weight data (Gusev et al. 2016) is available at http://gusevlab.org/projects/fusion/#reference-functional-data. An R package implementing our new methods is publicly available at https://github.com/yangq001/TEDE.

## 3 Results

### 3.1 Simulations

#### 3.1.1 Independent SNPs: Testing Horizontal Pleiotropy in MR

We generate genotype data of independent SNPs *G* = (*G_ij_*)_*n*×*p*_ using a multivariate binomial distribution, assuming Cov(*G_ij_, G_ik_*) = 0 (*j* ≠ *k*). We also assume each SNP has MAF *f* = 0.3 and simulate two traits X and Y using models similar to those in Slob et al. (2019):

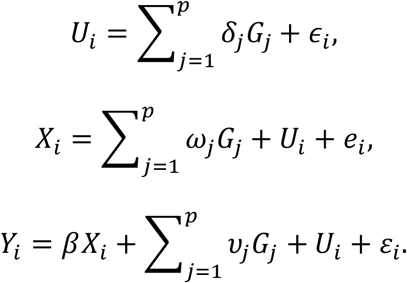

Here *ϵ_i_, e_i_* and *ε_i_* each follow an i.i.d standard normal distribution, *U_i_* is a confounder. *δ_j_, ω_j_, υ_j_* are the direct effects of SNP *j* on U, X and Y respectively. *β*, the causal effect of X on Y, is determined so that the proportion of variability in Y explained by X is about 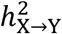. We generate *ω_j_*’s from a normal distribution with mean zero and standard deviation 0.15 first, and subsequently select those with *ω_j_* > 0.08 to avoid weak IVs, ensuring that the first valid IV assumption holds (i.e. an IV is associated with X). Then we shrink *ω_j_* by a constant so that the proportion of gene variance explained by SNPs is about 20%. We randomly choose some of the SNPs to be invalid IVs with horizontal pleiotropy, and we denote the proportion as %invalid. We set *υ_j_*’s as zero for valid IVs and as nonzero for invalid IVs. We consider the following different scenarios with a proportion (e.g. 0, 10%, 30%, 50%) of the IVs being invalid:

For invalid IVs, *υ_j_*~*N*(0,0.075). *δ_j_* = 0 for every IV. This means balanced pleiotropy. Here *α_j_* = *υ_j_*.
For invalid IVs, *υ_j_*~*N*(0.1 · sign(*ω_j_*), 0.025). *δ_j_* = 0 for every IV. This suggests directional pleiotropy. The direction of the direct effects is the same as that of G to X. Here *α_j_* = *υ_j_*.
For invalid IVs, *υ_j_~N*(0.1 · sign(*ω_j_*), 0.025). *δ_j_*~Unif(0,1) for invalid IVs. This suggests that, in addition to directional pleiotropy, the InSIDE (Instrument Strength Independent of Direct Effect) assumption is violated. Here *α_j_* = *υ_j_* + *δ_j_*. Note that (S1)(S2)(S3) actually become the same scenario when the proportion of invalid IVs is 0. In order to save space in the tables, we do not specify a different scenario for zero invalid IV.

When there are invalid IVs, we also shrink *υ_j_*’s so that the proportion of Y’s variance explained by 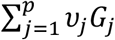 is about 0.3%. Note that choosing a higher proportion (e.g. 1%) will make the power of most tests much higher (e.g. very close to 1). Since we are looking at the two-sample setting, we generate one dataset with *n*_1_ subjects and another dataset with *n*_2_ subjects to obtain summary statistics for X and Y respectively. Then we apply different methods to the summary statistics and calculate their rejection rates based on 1000 simulations.

As shown in Table 1, when *p* = 30, all methods are able to control type I error rates with the default nominal significance level 0.05. All methods except MR-Egger have similar performance in both scenarios 1 and 2. MR-Egger has limited power even in the presence of directional pleiotropy, though its power increases as the proportion of invalid IVs goes up. TEDE-Sc’s power is higher than Cochran’s Q’s in all scenarios. TEDE-aSPU has higher power than TEDE-Sc when the proportion of invalid IVs is small, showing its advantage when dealing with sparse invalid IVs. In scenario 3, where the InSIDE assumption is violated in addition to directional pleiotropy, all methods have higher power, and the power goes up significantly as the proportion of invalid IVs increases. This power increase is different from what we have seen for scenarios 1 and 2 because in the first two scenarios, each invalid IV’s direct effect on Y (that does not go through X) is *α_j_* = *υ_j_*, but in the third scenario, it is *α_j_* = *υ_j_* + *υ_j_*. When the proportion of invalid IVs goes up, *υ_j_*’s tend to be smaller since we control the proportion of Y’s variance explained by 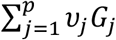, but *δ_j_*’s are not scaled, which means the total direct effect is much stronger with more invalid IVs in scenario 3, but not in scenarios 1 and 2.

**Table 1.** Rejection rates (type I error when there is 0 invalid IV; power otherwise) for testing horizontal pleiotropy. Independent variants. 1000 iterations. *p* = 30, *n*_1_ = 10000, *n*_2_ = 10000.

|  | 0 invalid | 10% invalid | 30% invalid | 50% invalid |
| --- | --- | --- | --- | --- |
| Scenario 1 (balanced pleiotropy), $\beta = 0$ | | | | |
| Cochran's Q | 0.037 | 0.848 | 0.853 | 0.832 |
| MR-Egger | 0.052 | 0.057 | 0.058 | 0.062 |
| TEDE-Sc | 0.045 | 0.859 | **0.874** | **0.856** |
| TEDE-aSPU | 0.039 | 0.927 | 0.859 | 0.8 |
| TEDE-Sc2 | 0.045 | 0.858 | 0.873 | 0.855 |
| TEDE-aSPU2 | 0.034 | **0.929** | 0.862 | 0.807 |
| Scenario 1 (balanced pleiotropy), $\beta > 0$ ($h_{X \rightarrow Y}^2 = 0.01$) | | | | |
| Cochran's Q | 0.041 | 0.803 | 0.812 | 0.782 |
| MR-Egger | 0.053 | 0.053 | 0.062 | 0.064 |
| TEDE-Sc | 0.048 | 0.821 | **0.829** | **0.803** |
| TEDE-aSPU | 0.039 | **0.891** | 0.816 | 0.756 |
| TEDE-Sc2 | 0.045 | 0.818 | 0.823 | 0.789 |
| TEDE-aSPU2 | 0.038 | 0.887 | 0.82 | 0.751 |
| Scenario 2 (directional pleiotropy), $\beta = 0$ | | | | |
| Cochran's Q | 0.037 | 0.843 | 0.827 | 0.757 |
| MR-Egger | 0.052 | 0.065 | 0.082 | 0.104 |
| TEDE-Sc | 0.045 | 0.856 | **0.855** | **0.773** |
| TEDE-aSPU | 0.039 | 0.928 | 0.811 | 0.729 |
| TEDE-Sc2 | 0.045 | 0.855 | **0.855** | 0.768 |
| TEDE-aSPU2 | 0.034 | **0.933** | 0.811 | 0.723 |
| Scenario 2 (directional pleiotropy), $\beta > 0$ ($h_{X \rightarrow Y}^2 = 0.01$) | | | | |
| Cochran's Q | 0.041 | 0.807 | 0.764 | 0.708 |
| MR-Egger | 0.053 | 0.06 | 0.094 | 0.117 |
| TEDE-Sc | 0.048 | 0.826 | **0.801** | **0.737** |
| TEDE-aSPU | 0.039 | **0.897** | 0.76 | 0.674 |
| TEDE-Sc2 | 0.045 | 0.815 | 0.785 | 0.722 |
| TEDE-aSPU2 | 0.038 | 0.894 | 0.747 | 0.661 |

| Scenario 3 (directional pleiotropy, InSIDE violated), $\beta = 0$ | | | | |
| --- | --- | --- | --- | --- |
| Cochran's Q | 0.037 | 0.826 | 0.971 | 0.99 |
| MR-Egger | 0.052 | 0.084 | 0.131 | 0.239 |
| TEDE-Sc | 0.045 | 0.839 | 0.984 | **1** |
| TEDE-aSPU | 0.039 | **0.853** | **0.987** | **1** |
| TEDE-Sc2 | 0.045 | 0.839 | 0.984 | **1** |
| TEDE-aSPU2 | 0.034 | 0.85 | 0.986 | **1** |
| Scenario 3 (directional pleiotropy, InSIDE violated), $\beta > 0$ ($h_{X \rightarrow Y}^2 = 0.01$) | | | | |
| Cochran's Q | 0.041 | 0.817 | 0.964 | 0.99 |
| MR-Egger | 0.053 | 0.077 | 0.122 | 0.219 |
| TEDE-Sc | 0.048 | 0.829 | **0.979** | **1** |
| TEDE-aSPU | 0.039 | **0.84** | 0.978 | **1** |
| TEDE-Sc2 | 0.045 | 0.825 | 0.978 | **1** |
| TEDE-aSPU2 | 0.038 | 0.834 | 0.976 | **1** |

Recall that, when there is no invalid IV, scenarios 1-3 all become the same (no invalid IVs; InSIDE not violated). This is why the type I error rates do not depend on different scenarios. MR-Egger depends on the InSIDE assumption and is usually expected to have problems with scenario 3, but that cannot be reflected in the type I errors here, since under the null hypothesis with no invalid IVs, InSIDE is always satisfied.

We further investigate the performance of each method with *p* = 100. As Table 2 shows, the power patterns in scenario 2 are similar to what we have in Table 1. We also have similar conclusions for scenario 3, the results of which are included in the supplementary materials (Table S1). TEDE-Sc seems to work better than Cochran’s Q, TEDE-aSPU works better when invalid IVs are sparse, and MR-Egger has low power. Nevertheless, when *β* > 0, Cochran’s Q, MR-Egger, TEDE-Sc and TEDE-aSPU have slightly inflated type I errors, and the inflation increases as *β* increases. This can be explained by looking at model 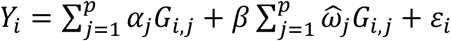. The true model under the null is 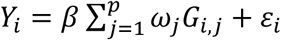, which means we should have 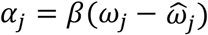. If the sample size for estimating 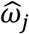’s is not large enough and *β* is nonzero, our estimate of *α_j_* can be off, which may lead to inflated type I errors. We need a sufficient sample size to ensure 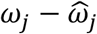 is small enough, especially when *p* is large. Cochran’s Q and MR-Egger may have similar issues even though their models are different. For instance, the direct effect in MR-Egger under the null is actually 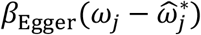, which means if 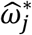’s are inaccurate and *β*_Egger_ is nonzero, the average of 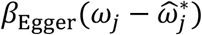’s may sometimes be nonzero and lead to inflated type I errors. As shown in Table 2, once we increase *n*_1_ to 50000, all methods are able to control type I errors. While usually the GWAS summary results used in MR analysis have sufficiently large samples (e.g. more than 100K), if we cannot obtain enough samples, we can use TEDE-Sc2 and TEDE-aSPU2, which are able to control type I errors better without losing much power as shown in Table 2.

**Table 2.** Rejection rates (type I error when there is 0 invalid IV; power otherwise) for testing horizontal pleiotropy. Independent variants. 1000 iterations. *p* = 100, *n*_1_ = 10000, *n*_2_ = 10000.

|  | 0 invalid | 10% invalid | 30% invalid | 50% invalid |
| --- | --- | --- | --- | --- |
| $n_1 = 10000, n_2 = 10000$ | | | | |
| Scenario 2 (directional pleiotropy), $\beta = 0$ | | | | |
| Cochran's Q | 0.048 | 0.559 | 0.501 | 0.49 |
| MR-Egger | 0.04 | 0.07 | 0.098 | 0.136 |
| TEDE-Sc | 0.043 | 0.578 | **0.521** | **0.532** |
| TEDE-aSPU | 0.046 | 0.672 | 0.45 | 0.425 |
| TEDE-Sc2 | 0.042 | 0.577 | 0.519 | 0.528 |
| TEDE-aSPU2 | 0.043 | **0.668** | 0.448 | 0.425 |
| Scenario 2 (directional pleiotropy), $\beta > 0$ ($h_{X \rightarrow Y}^2 = 0.02$) | | | | |
| Cochran's Q | 0.06 | 0.518 | 0.458 | 0.452 |
| MR-Egger | 0.071 | 0.126 | 0.182 | 0.242 |
| TEDE-Sc | 0.053 | 0.565 | **0.512** | **0.513** |
| TEDE-aSPU | 0.057 | **0.603** | 0.424 | 0.41 |
| TEDE-Sc2 | 0.048 | 0.512 | 0.454 | 0.453 |
| TEDE-aSPU2 | 0.049 | 0.574 | 0.383 | 0.353 |
| $n_1 = 50000, n_2 = 10000$ | | | | |
| Scenario 2 (directional pleiotropy), $\beta > 0$ ($h_{X \rightarrow Y}^2 = 0.02$) | | | | |
| Cochran's Q | 0.039 | 0.466 | 0.405 | 0.407 |
| MR-Egger | 0.055 | 0.067 | 0.109 | 0.118 |
| TEDE-Sc | 0.049 | 0.492 | **0.443** | **0.448** |
| TEDE-aSPU | 0.039 | **0.553** | 0.383 | 0.347 |
| TEDE-Sc2 | 0.044 | 0.486 | 0.429 | 0.432 |
| TEDE-aSPU2 | 0.043 | 0.549 | 0.376 | 0.339 |

#### 3.1.2 Correlated SNPs: Testing Horizontal Pleiotropy in TWAS

Now we generate genotype data of correlated SNPs *G* = (*G_ij_*)_*n*×*p*_. Following Barfield et al. (2017), we assume that the LD structure is AR(*ρ*) with Cov(*G_ij_, G_ik_*) = *ρ*^|*j*–*k*|^. We also assume each SNP has MAF *f* = 0.3. The rest is the same as what we did in the previous subsection for MR. Since we are looking at correlated variants, we only apply the LDA methods and examine their rejection rates based on 1000 simulations. Since *n*_1_ is usually relatively small in TWAS, we use *n*_1_ = 2000, *n*_2_ = 4000. Here we do not consider Cochran’s Q since it requires independent variants, and we replace MR-Egger with LDA MR-Egger. Furthermore, we include the recently developed PMR-Egger approach (Yuan et al. 2020) with its default setting, which can also use summary statistics of correlated SNPs to test horizontal pleiotropy.

As shown in Tables 3 and 4, when *p* = 30, most methods are able to control type I errors, while TEDE-Sc and TEDE-aSPU have much higher power than LDA MR-Egger in all scenarios. As expected, TEDE-Sc2 and TEDE-aSPU2 tend to be slightly more conservative than TEDE-Sc and TEDE-aSPU respectively. PMR-Egger has better performance than LDA MR-Egger when used to test horizontal pleiotropy in most cases, though its power is usually lower than that of TEDE. We also have similar findings for *p* = 100, for which more details are provided in the supplementary materials (Tables S2 and S3). Besides, we have observed some other interesting phenomena. For example, when the correlation between adjacent SNPs is relatively high, most methods seem to be more conservative in terms of smaller type I errors, and TEDE-aSPU has higher power than TEDE-Sc regardless of the proportion of invalid IVs, supporting its higher power for high-dimensional data. Further discussion on these is included in the supplementary materials as well.

**Table 3.** Rejection rates (type I error when there is 0 invalid IV; power otherwise) for testing valid IV assumptions. 1000 iterations. *p* = 30. Low LD (*ρ* = 0.3).

|  | 0 invalid | 10% invalid | 30% invalid | 50% invalid |
| --- | --- | --- | --- | --- |
| Scenario 1 (balanced pleiotropy), $\beta = 0$ | | | | |
| LDA MR-Egger | 0.05 | 0.103 | 0.093 | 0.1 |
| PMR-Egger | 0.043 | 0.135 | 0.145 | 0.134 |
| TEDE-Sc | 0.041 | 0.361 | **0.354** | **0.334** |
| TEDE-aSPU | 0.042 | **0.449** | 0.348 | 0.307 |
| TEDE-Sc2 | 0.041 | 0.36 | 0.351 | 0.329 |
| TEDE-aSPU2 | 0.046 | 0.447 | 0.346 | 0.297 |
| Scenario 1 (balanced pleiotropy), $\beta > 0$ ($h_{X \rightarrow Y}^2 = 0.01$) | | | | |
| LDA MR-Egger | 0.058 | 0.114 | 0.092 | 0.093 |
| PMR-Egger | 0.05 | 0.131 | 0.132 | 0.128 |
| TEDE-Sc | 0.049 | 0.345 | **0.343** | **0.328** |
| TEDE-aSPU | 0.048 | **0.407** | 0.332 | 0.284 |
| TEDE-Sc2 | 0.044 | 0.32 | 0.323 | 0.305 |
| TEDE-aSPU2 | 0.048 | 0.398 | 0.31 | 0.259 |
| Scenario 2 (directional pleiotropy), $\beta = 0$ | | | | |
| LDA MR-Egger | 0.05 | 0.111 | 0.105 | 0.111 |
| PMR-Egger | 0.043 | 0.146 | 0.137 | 0.106 |
| TEDE-Sc | 0.041 | 0.335 | **0.347** | **0.301** |
| TEDE-aSPU | 0.042 | **0.415** | 0.312 | 0.253 |
| TEDE-Sc2 | 0.041 | 0.328 | 0.338 | 0.294 |
| TEDE-aSPU2 | 0.046 | 0.42 | 0.309 | 0.246 |
| Scenario 2 (directional pleiotropy), $\beta > 0$ ($h_{X \rightarrow Y}^2 = 0.01$) | | | | |
| LDA MR-Egger | 0.058 | 0.141 | 0.145 | 0.154 |
| PMR-Egger | 0.05 | 0.138 | 0.125 | 0.106 |
| TEDE-Sc | 0.049 | 0.335 | **0.327** | **0.313** |
| TEDE-aSPU | 0.048 | **0.391** | 0.301 | 0.245 |
| TEDE-Sc2 | 0.044 | 0.307 | 0.29 | 0.274 |
| TEDE-aSPU2 | 0.048 | 0.377 | 0.281 | 0.212 |
| Scenario 3 (directional pleiotropy, InSIDE violated), $\beta = 0$ | | | | |
| LDA MR-Egger | 0.05 | 0.139 | 0.271 | 0.384 |
| PMR-Egger | 0.043 | 0.29 | 0.749 | 0.956 |
| TEDE-Sc | 0.041 | 0.587 | 0.866 | 0.968 |
| TEDE-aSPU | 0.042 | **0.646** | **0.885** | **0.973** |
| TEDE-Sc2 | 0.041 | 0.586 | 0.86 | 0.963 |
| TEDE-aSPU2 | 0.046 | 0.643 | 0.882 | 0.974 |
| Scenario 3 (directional pleiotropy, InSIDE violated), $\beta > 0$ ($h_{X \rightarrow Y}^2 = 0.01$) | | | | |
| LDA MR-Egger | 0.058 | 0.142 | 0.224 | 0.354 |
| PMR-Egger | 0.05 | 0.26 | 0.704 | 0.949 |
| TEDE-Sc | 0.049 | 0.566 | 0.855 | 0.962 |
| TEDE-aSPU | 0.048 | **0.618** | **0.867** | **0.97** |
| TEDE-Sc2 | 0.044 | 0.549 | 0.827 | 0.948 |
| TEDE-aSPU2 | 0.048 | 0.608 | 0.849 | 0.967 |

**Table 4.** Rejection rates (type I error when there is 0 invalid IV; power otherwise) for testing valid IV assumptions. 1000 iterations. *p* = 30. High LD (*ρ* = 0.7).

|  | 0 invalid | 10% invalid | 30% invalid | 50% invalid |
| --- | --- | --- | --- | --- |
| Scenario 1 (balanced pleiotropy), $\beta = 0$ | | | | |
| LDA MR-Egger | 0.046 | 0.164 | 0.135 | 0.149 |
| PMR-Egger | 0.056 | 0.25 | 0.299 | 0.265 |
| TEDE-Sc | 0.042 | 0.326 | 0.344 | 0.339 |
| TEDE-aSPU | 0.036 | **0.476** | **0.428** | **0.408** |
| TEDE-Sc2 | 0.041 | 0.325 | 0.341 | 0.335 |
| TEDE-aSPU2 | 0.036 | 0.47 | 0.423 | 0.402 |
| Scenario 1 (balanced pleiotropy), $\beta > 0$ ($h_{X \rightarrow Y}^2 = 0.01$) | | | | |
| LDA MR-Egger | 0.054 | 0.159 | 0.15 | 0.144 |
| PMR-Egger | 0.054 | 0.236 | 0.278 | 0.25 |
| TEDE-Sc | 0.052 | 0.319 | 0.33 | 0.313 |
| TEDE-aSPU | 0.04 | **0.449** | **0.404** | **0.385** |
| TEDE-Sc2 | 0.044 | 0.299 | 0.309 | 0.284 |
| TEDE-aSPU2 | 0.042 | 0.43 | 0.38 | 0.37 |

| Scenario 2 (directional pleiotropy), $\beta = 0$ | | | | |
| --- | --- | --- | --- | --- |
| LDA MR-Egger | 0.046 | 0.161 | 0.145 | 0.154 |
| PMR-Egger | 0.056 | 0.24 | 0.289 | 0.231 |
| TEDE-Sc | 0.042 | 0.34 | 0.332 | 0.276 |
| TEDE-aSPU | 0.036 | **0.464** | **0.389** | **0.348** |
| TEDE-Sc2 | 0.041 | 0.338 | 0.325 | 0.267 |
| TEDE-aSPU2 | 0.036 | 0.462 | 0.393 | 0.347 |
| Scenario 2 (directional pleiotropy), $\beta > 0$ ($h_{X \rightarrow Y}^2 = 0.01$) | | | | |
| LDA MR-Egger | 0.054 | 0.173 | 0.17 | 0.166 |
| PMR-Egger | 0.054 | 0.225 | 0.253 | 0.222 |
| TEDE-Sc | 0.052 | 0.329 | 0.331 | 0.289 |
| TEDE-aSPU | 0.04 | **0.439** | **0.382** | **0.332** |
| TEDE-Sc2 | 0.044 | 0.297 | 0.296 | 0.25 |
| TEDE-aSPU2 | 0.042 | 0.424 | 0.364 | 0.317 |
| Scenario 3 (directional pleiotropy, InSIDE violated), $\beta = 0$ | | | | |
| LDA MR-Egger | 0.046 | 0.23 | 0.438 | 0.625 |
| PMR-Egger | 0.056 | 0.476 | 0.907 | 0.989 |
| TEDE-Sc | 0.042 | 0.576 | 0.905 | 0.986 |
| TEDE-aSPU | 0.036 | **0.656** | 0.939 | **0.99** |
| TEDE-Sc2 | 0.041 | 0.569 | 0.896 | 0.982 |
| TEDE-aSPU2 | 0.036 | 0.649 | **0.94** | 0.987 |
| Scenario 3 (directional pleiotropy, InSIDE violated), $\beta > 0$ ($h_{X \rightarrow Y}^2 = 0.01$) | | | | |
| LDA MR-Egger | 0.054 | 0.219 | 0.417 | 0.615 |
| PMR-Egger | 0.054 | 0.459 | 0.895 | 0.987 |
| TEDE-Sc | 0.052 | 0.559 | 0.895 | 0.985 |
| TEDE-aSPU | 0.04 | **0.634** | **0.93** | **0.99** |
| TEDE-Sc2 | 0.044 | 0.532 | 0.876 | 0.978 |
| TEDE-aSPU2 | 0.042 | 0.615 | 0.923 | 0.983 |

### 3.2 Real Data Applications

#### 3.2.1 Testing Direct Effects in MR and TWAS for SCZ and Other Complex Traits

Richardson et al. (2019) found some strong evidence for causal relationships between genetic liability to SCZ (schizophrenia) and many complex traits by constructing polygenic risk scores (PRS) for association analyses. Further investigations were done with a two-sample MR analysis to back up the conclusions. We apply different methods to test for direct effects in a similar context to see whether there are noticeable invalid IVs that may cast doubts on the conclusions of the MR analysis of SCZ and the complex traits. For SCZ, we use a GWAS summary dataset based on 150K subjects from the Schizophrenia Working Group of the Psychiatric Genomics Consortium (2014). As for other complex traits, we choose eight of the traits included in Richardson et al. (2019)’s MR analysis. For these traits (listed in Table 5), we use the GWAS results based on the imputed UK Biobank data (Sudlow et al. 2015; Neale Lab 2017) with up to 362K subjects. This is a two-sample problem since the subjects do not overlap. We check out the SNPs whose minor allele frequencies are greater than 0.1 and whose p-values for marginal associations with SCZ are smaller than 5e-8. Then we select those that are also present in the 1000 Genomes Project Data (phase 3; 503 subjects with European ancestry) from the 1000 Genomes Project Consortium (2012). We prune the SNPs based on the LD information estimated from the 1000 Genomes data to get independent SNPs (*r*^2^ < 0.001), resulting in 39 IVs selected for MR analysis. We apply both the non-LDA tests and the LDA tests to test for direct effects with Y being each of the complex traits of interest.

**Table 5.** P-values of testing direct effects for selected IVs. Exposure: SCZ. Significance threshold: 5e-3. TEDE-aSPU and TEDE-aSPU2 use 1e+4 iterations.

| 39 independent IVs |  |  |  |  |  |  |
| --- | --- | --- | --- | --- | --- | --- |
|  | Cochran's Q | MR-Egger | TEDE-Sc | TEDE-Sc2 | TEDE-aSPU | TEDE-aSPU2 |
| Tense* | 0 | 1.4e-1 | 0 | 0 | <1e-4 | <1e-4 |
| Psychiatrist* | 3.0e-13 | 8.0e-3 | 1.2e-13 | 1.2e-13 | <1e-4 | <1e-4 |
| Depression* | 2.5e-3 | 1.8e-2 | 1.9e-3 | 1.9e-3 | 8.0e-3 | 8.5e-3 |
| Neuroticism* | 0 | 2.3e-1 | 0 | 0 | <1e-4 | <1e-4 |
| Fluid Int* | 0 | 3.2e-1 | 0 | 0 | <1e-4 | <1e-4 |
| Matches* | 0 | 1.1e-1 | 0 | 0 | <1e-4 | <1e-4 |
| Stop-Smoking* | 3.7e-11 | 4.4e-2 | 1.6e-11 | 1.6e-11 | <1e-4 | <1e-4 |
| Past Smoking* | 0 | 7.0e-1 | 0 | 0 | <1e-4 | <1e-4 |
| 140 correlated IVs |  |  |  |  |  |  |
|  | LDA MR-Egger | TEDE-Sc | TEDE-Sc2 | TEDE-aSPU | TEDE-aSPU2 |  |
| Tense* | 6.3e-3 | 0 | 0 | <1e-4 | <1e-4 |  |
| Psychiatrist* | 9.8e-3 | 0 | 0 | <1e-4 | <1e-4 |  |
| Depression* | 1.4e-2 | 0 | 0 | <1e-4 | <1e-4 |  |
| Neuroticism* | 3.6e-2 | 0 | 0 | <1e-4 | <1e-4 |  |
| Fluid Int* | 7.5e-1 | 0 | 0 | <1e-4 | <1e-4 |  |
| Matches* | 2.8e-3 | 0 | 0 | <1e-4 | <1e-4 |  |
| Stop-Smoking* | 1.7e-2 | 0 | 0 | <1e-4 | <1e-4 |  |
| Past Smoking* | 1.7e-1 | 0 | 0 | <1e-4 | <1e-4 |  |

\*Abbreviations for Tense / 'highly strung'; Seen a psychiatrist for nerves, anxiety, tension or depression; Non-cancer illness code: self-reported: depression; Neuroticism score; Fluid intelligence score; Number of incorrect matches in round; Number of unsuccessful stop-smoking attempts; Past tobacco smoking.

As shown in Table 5, with 39 independent IVs, most of the tests have highly significant results for most of the analyzed outcomes. MR-Egger does not give any significant p-values under level 5e-3, which is consistent with its low power shown in our simulation studies, especially when the pleiotropy is not directional. TEDE-Sc’s p-values are usually smaller than those of Cochran’s Q, which is also consistent with our previous finding that TEDE-Sc tends to have higher power than Cochran’s Q. TEDE-Sc2 and TEDE-aSPU2 are very close to TEDE-Sc and TEDE-aSPU, probably because 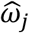’s have very small standard deviations given the large sample size for SCZ.

For this problem, TWAS with GWAS summary statistics can also be applied to examine the relationship between SCZ and other traits, which may be more powerful by including more and correlated SNPs as IVs. As in MR, we need to test for direct effects as a way to check whether the TWAS model is appropriate. We use the LDA methods to test direct effects for each exposure-outcome pair using correlated SNPs across different chromosomes. This time we select significant SNPs based on *r*^2^ < 0.025 instead of *r*^2^ < 0.001, resulting in 140 SNPs. As shown in Table 5, TEDE-Sc and TEDE-aSPU have highly significant results, while LDA MR-Egger does not. For self-reported depression, the p-value is much more significant than before after including correlated SNPs, which may confirm the potential downside of adding more SNPs as IVs. Including more correlated IVs may yield higher power, but at the same time it will increase the chance of having invalid IVs.

#### 3.2.2 Testing Direct Effects in TWAS of AD

TWAS can be very useful in identifying genes whose expression contributes to complex traits like Alzheimer’s disease (AD). By making use of correlated SNPs, instead of only independent SNPs, TWAS (as a more general MR approach) can be more powerful than MR as shown in certain scenarios (Knutson and Pan 2020). However, similar to what MR faces, TWAS may gave incorrect results due to the violation of the valid IV assumptions, including the assumption of no horizontal pleiotropy. We use the ADNI data (Shen et al. 2014), the IGAP stage 1 data (Lambert et al. 2013) and the reference gene expression weights from Gusev et al. (2016), which we call the weight data, to test whether direct effects of IVs exist in TWAS analysis of each gene and AD.

The weight data contains 6007 genes’ expression in the whole blood. For each gene’s expression, the weight data provides the SNPs selected by the elastic net regression and their joint effect sizes on the gene’s expression, which we use as our 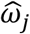’s. These were pre-computed based on 369 samples. The IGAP GWAS summary statistics data (with a sample size of 54162) are used along with the ADNI individual-level data (sample size 712) to calculate 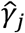’s and their covariance matrix. We match the SNPs across different datasets and prune them to make sure none of the pairwise correlations is larger than 0.9 (in absolute values). For convenience and to be consistent with the previous sections, we define each locus as the SNPs selected for each gene expression. We exclude those loci with less than 5 SNPs, resulting in 3611 loci and 49225 SNPs remaining. Next, we apply the various tests using GWAS summary statistics to test for direct effects for each locus. Since the number of loci is large and the significance threshold is very small, we choose to use the asymptotics-based TEDE-aSPU to save computation time. TEDE-Sc2, TEDE-aSPU2 and PMR-Egger cannot be applied since the variance of 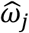 is not provided in the weight data.

As Figure 2 (A) shows, many loci have been detected to have direct effects, suggesting that many SNPs may affect AD through pathways other than the corresponding gene. TEDE-Sc and TEDE-aSPU have found many more significant loci than LDA MR-Egger. TEDE-Sc is able to detect more loci with horizontal pleiotropy than TEDE-aSPU, probably suggesting that the proportion of invalid IVs is usually relatively high. Meanwhile, some loci only appear to be significant according to TEDE-aSPU, showing the complementary role of the two versions of the TEDE test.

**Figure 2.**
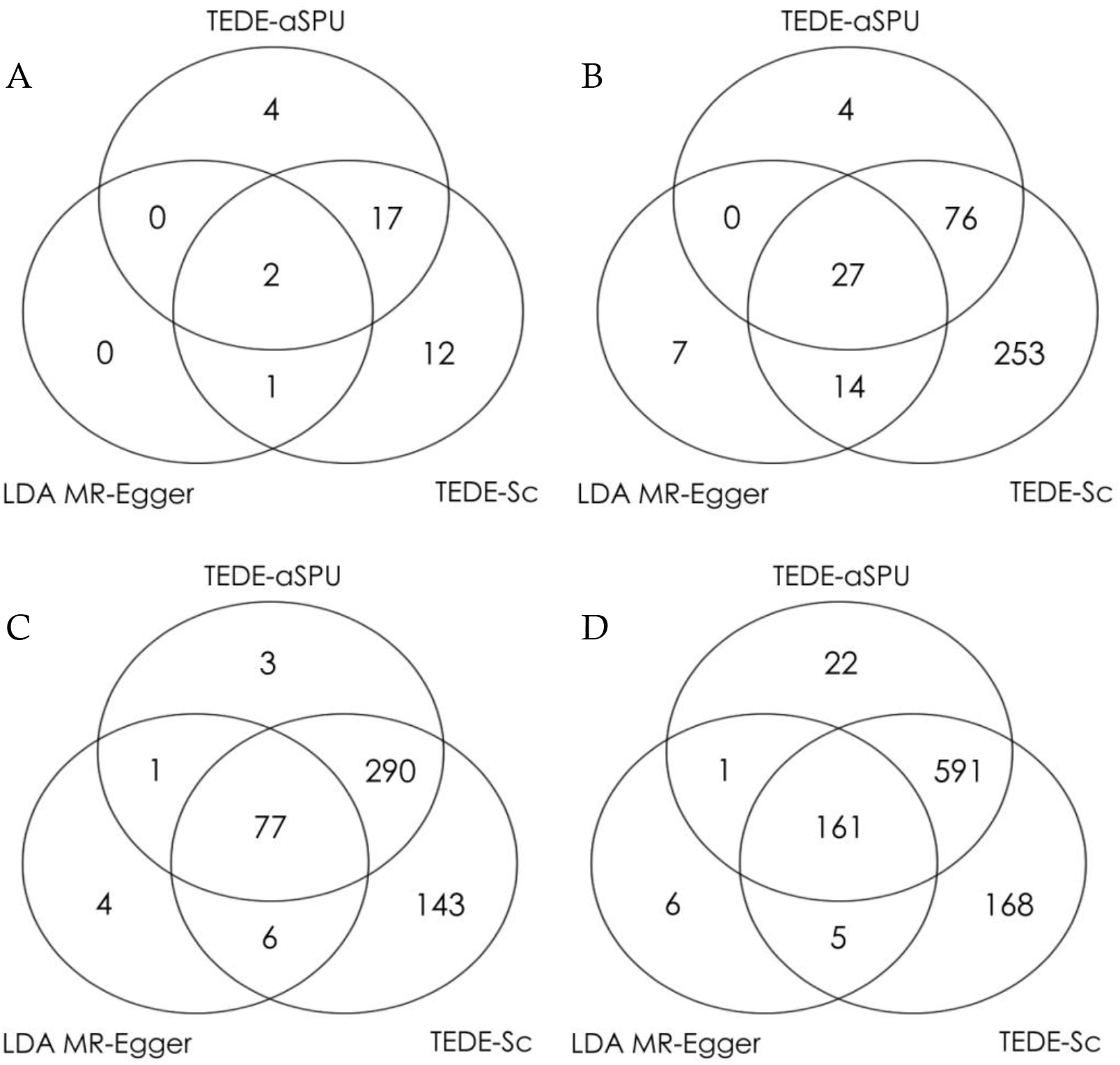
Numbers of significant loci with direct effects. Significance threshold: 0.05/#loci. A: AD as outcome (3611 loci with 49225 SNPs). B: SCZ as outcome (3611 loci with 49225 SNPs). C: LDL as outcome (2010 lipid data; 4267 loci with 58382 SNPs). D: LDL as outcome (2013 lipid data; 4267 loci with 58382 SNPs).

Compared to the total number of loci, the proportion of loci with detected direct effects may seem small. However, it makes sense because most of the SNPs and genes cannot be detected to be associated with AD. In such a case, we do expect that direct effects of SNP to AD are not detectable or even do not exist for most loci. However, we still need to be careful when we have significant TWAS results. After applying TWAS and LDA MR-Egger to test the association between each gene and AD, we list the significant loci in Table 6 along with the p-values of testing direct effects. TEDE-Sc has detected direct effects in three of the seven loci, covering the two found by TEDE-aSPU and the two found by LDA MR-Egger. Also, if we examine the 21 loci with at least one SNP’s marginal p-value smaller than 5e-6 for its association with AD, 9.5%, 43% and 48% of them are found to have direct effects by LDA MR-Egger, TEDE-Sc and TEDE-aSPU respectively under the same significance threshold 1.38e-5. These results demonstrate the power advantage of TEDE-Sc and TEDE-aSPU over LDA MR-Egger, as well as the need to test for direct effects for model checking in TWAS, especially when we obtain significant associations.

**Table 6.** P-values of testing gene expression to AD effects and other direct effects. 3611 loci (49225 SNPs) were tested in total. Stars indicate achieving statistical significance at Bonferroni adjusted significance threshold 1.38e-5.

| Chr | Gene | Testing gene expression to AD effects |  | Testing direct effects |  |  |
| --- | --- | --- | --- | --- | --- | --- |
|  |  | TWAS | LDA MR-Egger | LDA MR-Egger | TEDE-Sc | TEDE-aSPU |
| 1 | PTGFR | 2.7e-4 | **4.8e-7\*** | 4.5e-4 | 2.0e-2 | 1.6e-1 |
| 2 | MTG1 | **5.3e-6\*** | 8.9e-3 | 5.5e-1 | **7.9e-9\*** | **1.5e-10\*** |
| 7 | MIS12 | 5.4e-3 | **6.0e-6\*** | 3.6e-4 | 8.3e-3 | 9.8e-5 |
| 7 | GRAP | **6.5e-10\*** | 1.1e-4 | 1.0e-1 | 8.6e-4 | 3.0e-2 |
| 11 | MITD1 | 6.3e-2 | **2.7e-7\*** | **1.5e-6\*** | **2.6e-6\*** | 1.7e-5 |
| 11 | CAPN13 | **2.7e-6\*** | 5.1e-2 | 6.3e-1 | 7.5e-3 | 8.5e-2 |
| 19 | POMZP3 | 1.3e-1 | **1.2e-24\*** | **2.1e-57\*** | **0\*** | **0\*** |

#### 3.2.3 Testing Direct Effects in TWAS of SCZ and LDL

To further examine the existence of horizontal pleiotropy in other scenarios, we apply TWAS for SCZ with the SCZ data used in the previous section and for LDL (low-density lipoprotein cholesterol) with the 2010 and 2013 lipid data (Teslovich et al. 2010; Willer et al. 2013) separately. As shown in Figure 2 (B, C, D), after similar analyses, we are able to detect many significant loci with direct effects, especially for LDL: about 20% of the loci are significant. This is consistent with our previous explanation since more SNPs are associated with LDL than SCZ and AD. We also find that using the 2013 lipid data gives more significant results than using the 2010 lipid data, probably because of the sample size difference (about 189K vs. 100K). These results further demonstrate the possibility of widespread horizontal pleiotropy and the need for model checking in TWAS.

## 4 Discussion

We have presented a novel method with two versions (TEDE-Sc and TEDE-aSPU) that can be applied to test for direct effects as a general GOF test for model checking in MR and TWAS. For MR with only independent IVs across different loci, our simulations show that TEDE-Sc is more powerful than the widely used Cochran’s Q statistic in most cases. TEDE-aSPU works better than TEDE-Sc when the proportion of invalid IVs is small and/or the number of invalid IVs is large. MR-Egger has quite limited power when compared to other methods even in the presence of strong directional pleiotropy. We have noticed that when the number of IVs is large (e.g. ~100) and the sample size for the exposure is not large enough, the tests with higher power may have slightly inflated type I errors (for detecting horizontal pleiotropy). Our alternative versions of the new method, TEDE-Sc2 and TEDE-aSPU2, are able to control type I errors better by taking into account the variability of estimating the effects of the SNPs/IVs on the exposure; however, these two versions require the summary level data to contain the standard errors of the estimated effects of SNPs on the exposure/X, (i.e. se(ωj)’s or se(?y)’s). After applying different methods to test for direct effects in an MR analysis of SCZ and some complex traits, almost all of the results from Cochran’s Q, TEDE-Sc and TEDE-aSPU turned out to be significant, indicating that the conclusions from the MR analysis may be problematic given the strong evidence of wide-spread direct effects. Meanwhile, MR-Egger did not reject the null hypothesis, presumably due to its low power or the possibility that the pleiotropy is not directional or uncorrelated (i.e. when the InSIDE assumption does not hold), confirming the potential issue of using MR-Egger (or its modification like LDA MR-Egger) for model checking in MR (or TWAS) in spite of its wide use in practice (Bowden et al 2018).

For TWAS, which usually includes correlated SNPs associated with a gene’s expression level, TEDE-Sc and TEDE-aSPU are able to make use of the LD information and control type I errors like LDA MR-Egger. Nevertheless, similar to MR-Egger, LDA MR-Egger is fairly low powered when used to test for horizontal pleiotropy in our simulations. Again, TEDE-Sc seems to work better when the proportion of invalid IVs is relatively high, while TEDE-aSPU can handle the sparse or high-dimensional situation better. When different LDA methods were applied to test for horizontal pleiotropy in a TWAS analysis of gene expression and AD, TEDE-Sc identified many significant loci, while TEDE-aSPU found much fewer, which might suggest that in many of these loci there were a relatively high proportion of SNPs with horizontal pleiotropy. On the other hand, TEDE-aSPU managed to find some significant loci that were not detected by TEDE-Sc, showing their complementary roles. In another real data application, our new method found around 20% of the loci with horizontal pleiotropy in TWAS of LDL using a large-scale GWAS lipid data, demonstrating substantial power advantages of our new tests over LDA MR-Egger and more importantly, highlighting the need to test for horizontal pleiotropy as a way to check the TWAS modeling assumptions in practice.

In practice, if it is reasonable to assume sparse direct effects, especially with a relatively large number of the SNPs/IVs to be tested, we’d recommend TEDE-aSPU2 for its expected higher power; otherwise, we recommend TEDE-Sc2 for its generally better performance as shown in our data examples (Figure 2) and often a relatively small number of the SNPs/IVs being tested. Alternatively, one may apply both TEDE-Sc2 and TEDE-aSPU2 to detect direct effects, given their different power advantages in different scenarios while controlling type I errors well. However, in certain situations where the summary level data of the exposure only have joint SNP-to-exposure effect size estimates 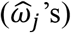 but without their standard errors, the above two cannot be applied and, instead, TEDE-Sc and TEDE-aSPU can be applied. We would also like to point out that our methods run reasonably fast with reasonable numbers of IVs: Table S3 in the supplementary materials contains more information on their computing time.

In the future, we can extend our new method to other MR or TWAS applications, such as MV-TWAS (Knutson et al. 2020), which is a more robust version of TWAS (or MR) by including multiple genes (or other traits) as multiple exposures in the same model. In this scenario, we can test whether there are direct effects of the SNPs on the outcome through pathways other than through any of the multiple exposures included in the model; lack of any evidence in such a test would lend support for the goodness-of-fit of the MV-TWAS model, and thus support for its conclusions. Besides, as suggested by a reviewer, identifying then removing invalid IVs in an MR analysis may lead to better results (Xue et al 2021). Our current methods aim for global testing (on whether there is any direct effect by any IV), rather than testing on each individual IVs. The latter task may appear straightforward to implement based on our framework, but it may be challenging to accurately identify which IVs are invalid because of the difficulty in accurately estimating each direct effect in the over-specified model (1) (in which the parameters are non-identifiable if all SNPs/IVs used in imputing X are included for their possible direct effects on Y). On the other hand, our proposed methods are based on the simplified and identifiable model under the null hypothesis. Nevertheless, it is possible to develop a sequential testing procedure to filter out invalid IVs one at a time as shown in a different approach of Xue et al (2021), or other penalized regression and variable selection methods (Windmeijer et al. 2016, 2019); this is worth further investigation.

## Acknowledgements

We thank the reviewers for many insightful and helpful comments. This research was supported by NIH grants RF1AG067924, R01AG065636, R01HL116720, R01GM113250 and R01GM126002, by NSF grant DMS 1711226, and by the Minnesota Supercomputing Institute at the University of Minnesota.

The ADNI data collection and sharing for this project was funded by the Alzheimer’s Disease Neuroimaging Initiative (ADNI) (National Institutes of Health Grant U01 AG024904) and DOD ADNI (Department of Defense award number W81XWH-12-2-0012). ADNI is funded by the National Institute on Aging, the National Institute of Biomedical Imaging and Bioengineering, and through generous contributions from many other institutes.

## List of Support Information Legends

Additional Explanations of Tables 3 and 4 (Main Text)

Additional Explanation to that Population Structure Can Lead to Some Direct Effects of SNPs/IVs

### Additional Figures

**Figure S1.** Flowchart for common MR/TWAS analysis with summary level data.

**Figure S2.** Two different scenarios of having direct effects or “correlated pleiotropy”: (A) G affects U. (B) U affects G.

### Additional Tables

**Table S1.** Rejection rates (type I error when there is 0 invalid IV; power otherwise) for testing horizontal pleiotropy. Independent variants. 1000 iterations. *p* = 100, *n*_1_ = 10000, *n*_2_ = 10000.

**Table S2.** Rejection rates (type I error when there is 0 invalid IV; power otherwise) for testing valid IV assumptions. 1000 iterations. *p* = 100. Low LD (*ρ* = 0.3).

**Table S3.** Rejection rates (type I error when there is 0 invalid IV; power otherwise) for testing valid IV assumptions. 1000 iterations. *p* = 100. High LD (*ρ* = 0.7).

**Table S4.** Computation time of TEDE (seconds) averaged over 5 runs. TEDE-Sc gives the results for both TEDE-Sc and TEDE-Sc2, and TEDE-aSPU gives the results for both TEDE-aSPU and TEDE-aSPU2.

**Table S5.** Summary of the datasets used in the manuscript.

