## Supplemental Materials for "Model Checking via Testing for Direct Effects in Mendelian Randomization and Transcriptome-wide Association Studies"

#### Additional Explanations of Tables 3 and 4 (Main Text)

Based on Tables 3 and 4 in the main text, when the correlation between adjacent variants is relatively high, most methods seem to be more conservative in terms of type I errors. This can be partially explained by the same argument used in the previous section (3.1.1, main text) that the direct effect under the null is actually $\alpha_j = \beta(\omega_j - \widehat{\omega}_j)$. Based on our experience, if $\widehat{\omega}_j$'s are very accurate (almost the same as $\omega_j$), the type I error rate is usually controlled at 0.05. Nevertheless, when they are not that accurate, we may not get the ideal type I error rate. In general, if the sample sizes are fixed, increasing $\beta$ and increasing $\omega_j$ are both likely to increase type I errors. Since we control the variance of $X_i$ explained by $\sum_{j=1}^{p} \omega_j G_j$ to be 0.2, when the pairwise correlation between adjacent SNPs is positive, the larger this correlation is, the smaller $\omega_j$'s tend to be, which is why the type I error rates in Table 5 tend to be smaller than those in Table 4. We also find through some additional experiments that if we increase the proportion of the variance explained (e.g. from 0.2 to 0.5), the type I errors tend to go up. Since in practice we do not know whether $\beta$ and $\omega_j$'s are large enough to create inflated type I errors, it is better to use a larger sample and apply methods like TEDE-Sc2 and TEDE-aSPU2 whenever possible.

Another interesting phenomenon is that with a relatively high LD correlation, TEDE-aSPU has higher power than TEDE-Sc regardless of the proportion of invalid IVs. This may be a result of their test statistics using the LD matrix differently. TEDE-Sc uses $\mathbf{U}(\hat{\boldsymbol{\theta}}_0)'\mathbf{I}^{-1}(\hat{\boldsymbol{\theta}}_0)\mathbf{U}(\hat{\boldsymbol{\theta}}_0)$, which is like transforming the scores to make them uncorrelated and then taking the sum of squares. TEDE-aSPU simply takes the sum of powered scores without decorrelation, which may be more efficient when the SNPs are positively correlated than uncorrelated.

#### Additional Explanation to that Population Structure Can Lead to Some Direct Effects of SNPs/IVs

In addition to invalid IVs, there are other reasons for the violation of modeling assumptions in MR and TWAS. Here we show that, like invalid IVs, population structure can lead to some direct effects (appearing as "correlated pleiotropy") of SNPs/IVs on the outcome, to which our proposed TEDE test can be applied.

For correlated pleiotropy with effects between IVs and confounders, there are two possibilities, illustrated in Figure S2. It is easy to see that when there is a causal effect from G (as IVs) to U (Figure S2 (A)), it will lead to horizontal pleiotropy. We will demonstrate that when there is an effect from U to G (Figure S2 (B)), it will also lead to some direct effects appearing as "horizontal pleiotropy". In this scenario, we assume

$$\mathbf{G} = \mathbf{U}\boldsymbol{\varsigma} + \mathbf{e}_G,$$

where $\mathbf{U}$ is a $n$ by $l$ matrix denoting $l$ confounders for $n$ subjects, $\boldsymbol{\varsigma}$ is a $l$ by $p$ matrix denoting the effects of $l$ confounders on $p$ IVs. $\mathbf{e}_G$ denotes the part of $\mathbf{G}$ not explained by $\mathbf{U}$. This means

$$\mathbf{U} = \boldsymbol{\varsigma}^{-1}\mathbf{G} + \boldsymbol{\varsigma}^{-1}\mathbf{e}_G,$$

$$\mathrm{E}(\mathbf{U}|\mathbf{G}) = \boldsymbol{\varsigma}^{-1}\mathbf{G} + \boldsymbol{\varsigma}^{-1}\mathrm{E}(\mathbf{e}_G|\mathbf{G}).$$

Given that $\mathbf{X} = \mathbf{G}\boldsymbol{\omega} + \mathbf{U}\boldsymbol{\phi} + \mathbf{e}_X$, and assuming $\mathbf{e}_X$ has mean 0 since the exposure has been centered at 0, we have

$$\mathrm{E}(\mathbf{X}|\mathbf{G}) = \mathbf{G}\boldsymbol{\omega} + \boldsymbol{\varsigma}^{-1}\mathbf{G}\boldsymbol{\phi} + \boldsymbol{\varsigma}^{-1}\mathrm{E}(\mathbf{e}_G|\mathbf{G})\boldsymbol{\phi}.$$

Since $\mathbf{Y} = \beta\mathbf{X} + \mathbf{U}\boldsymbol{\zeta} + \mathbf{e}_Y$ with $\mathbf{e}_Y$ centered at 0, we have

$$\mathrm{E}(\mathbf{Y}|\mathbf{G}) = \beta \mathrm{E}(\mathbf{X}|\mathbf{G}) + \boldsymbol{\varsigma}^{-1}\mathbf{G}\boldsymbol{\zeta} + \boldsymbol{\varsigma}^{-1}\mathrm{E}(\mathbf{e}_G|\mathbf{G})\boldsymbol{\zeta}.$$

Hence, we have $\mathbf{G}$'s effects on Y not only through $\mathbf{X}$ (i.e. shown by the term $\mathrm{E}(\mathbf{X}|\mathbf{G})$) but also on Y directly (i.e. through $\boldsymbol{\varsigma}^{-1}\mathbf{G}\boldsymbol{\zeta} + \boldsymbol{\varsigma}^{-1}\mathbf{E}(\mathbf{e}_G|\mathbf{G})\boldsymbol{\zeta}$), which are the direct effects manifested as "horizontal pleiotropy". As a result, testing the null hypothesis of no direct effects using TEDE can help us detect the violation modeling assumptions (e.g. in the presence of population structure) in the scenario illustrated in Figure S2(B) as well.

Note that we use a linear model, instead of a logistic regression model, in modeling the effects of population structure $\mathbf{U}$ on $\mathbf{G}$ for the following two reasons. First, since we usually use some common variants in $\mathbf{G}$ with their minor/major allele frequencies bounded away from 0 and ½, a linear model is reasonable and general. $\mathbf{U}$ can be categorical, indicating different subpopulations, or be continuous, representing the underlying population structure, e.g. as some top principle components. Second, the use of the linear model largely simplifies the above derivation to illustrate our point.

### Additional Figures

**Figure S1.** Flowchart for common MR/TWAS analysis with summary level data.

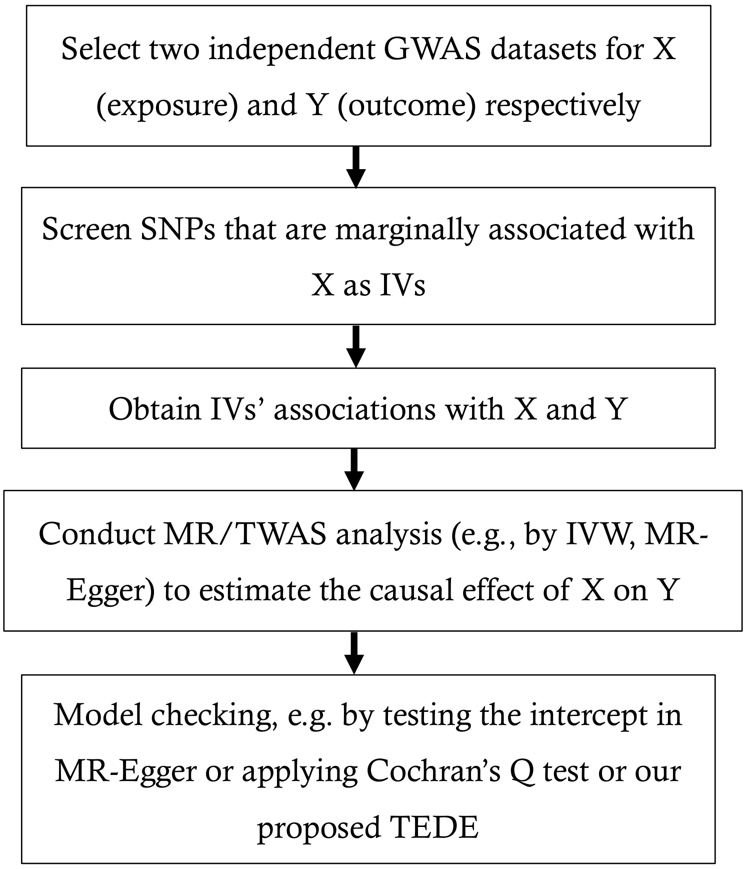


**Figure S2.** Two different scenarios of having direct effects or "correlated pleiotropy": (A) G affects U. (B) U affects G.

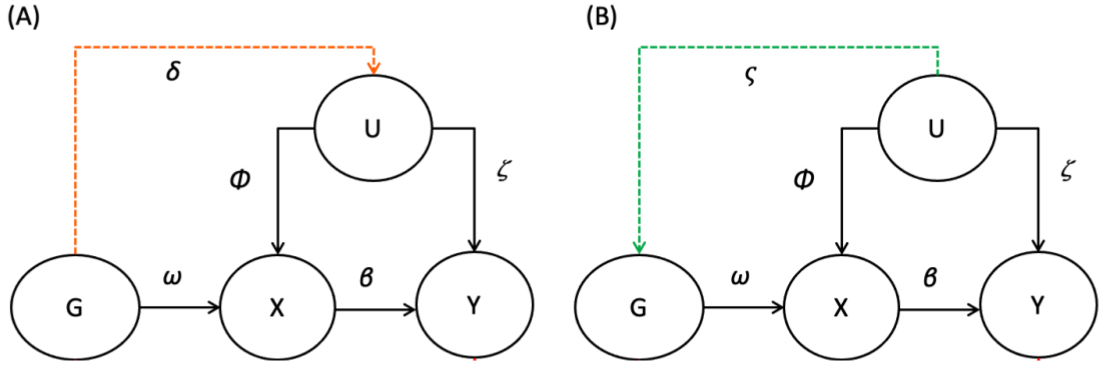

### Additional Tables

**Table S1.** Rejection rates (type I error when there is 0 invalid IV; power otherwise) for testing horizontal pleiotropy. Independent variants. 1000 iterations. $p = 100, n_1 = 10000, n_2 = 10000$.

| | 0 invalid | 10% invalid | 30% invalid | 50% invalid |
|---|---|---|---|---|
| $n_1 = 10000, n_2 = 10000$ | | | | |
| Scenario 3 (directional pleiotropy), $\beta = 0$ | | | | |
| Cochran's Q | 0.048 | 0.958 | 0.982 | 0.979 |
| MR-Egger | 0.04 | 0.148 | 0.371 | 0.489 |
| TEDE-Sc | 0.043 | 0.98 | 1 | 1 |
| TEDE-aSPU | 0.046 | 0.989 | 1 | 1 |
| TEDE-Sc2 | 0.042 | 0.98 | 1 | 1 |
| TEDE-aSPU2 | 0.043 | 0.988 | 1 | 1 |
| Scenario 3 (directional pleiotropy), $\beta > 0$ ($h^2_{\mathrm{X}\to\mathrm{Y}} = 0.02$) | | | | |
| Cochran's Q | 0.06 | 0.95 | 0.982 | 0.979 |
| MR-Egger | 0.071 | 0.09 | 0.288 | 0.411 |
| TEDE-Sc | 0.053 | 0.972 | 1 | 1 |
| TEDE-aSPU | 0.057 | 0.985 | 1 | 1 |
| TEDE-Sc2 | 0.048 | 0.966 | 1 | 1 |
| TEDE-aSPU2 | 0.049 | 0.98 | 1 | 1 |
| $n_1 = 50000, n_2 = 10000$ | | | | |
| Scenario 3 (directional pleiotropy), $\beta > 0$ ($h^2_{\mathrm{X}\to\mathrm{Y}} = 0.02$) | | | | |
| Cochran's Q | 0.039 | 0.947 | 0.971 | 0.978 |
| MR-Egger | 0.055 | 0.178 | 0.432 | 0.572 |
| TEDE-Sc | 0.049 | 0.963 | 1 | 1 |
| TEDE-aSPU | 0.039 | 0.977 | 1 | 1 |
| TEDE-Sc2 | 0.044 | 0.962 | 1 | 1 |
| TEDE-aSPU2 | 0.043 | 0.973 | 1 | 1 |

**Table S2.** Rejection rates (type I error when there is 0 invalid IV; power otherwise) for testing valid IV assumptions. 1000 iterations. $p = 100$. Low LD ($\rho = 0.3$).

| | 0 invalid | 10% invalid | 30% invalid | 50% invalid |
|---|---|---|---|---|
| Scenario 1 (balanced pleiotropy), $\beta = 0$ | | | | |
| LDA MR-Egger | 0.051 | 0.076 | 0.059 | 0.065 |
| PMR-Egger | 0.062 | 0.08 | 0.073 | 0.068 |
| TEDE-Sc | 0.038 | 0.168 | 0.174 | 0.163 |
| TEDE-aSPU | 0.04 | 0.205 | 0.167 | 0.146 |
| TEDE-Sc2 | 0.036 | 0.168 | 0.169 | 0.158 |
| TEDE-aSPU2 | 0.04 | 0.199 | 0.158 | 0.134 |
| Scenario 1 (balanced pleiotropy), $\beta > 0$ ($h^2_{X\to Y} = 0.01$) | | | | |
| LDA MR-Egger | 0.065 | 0.081 | 0.07 | 0.077 |
| PMR-Egger | 0.057 | 0.072 | 0.07 | 0.069 |
| TEDE-Sc | 0.049 | 0.177 | 0.191 | 0.161 |
| TEDE-aSPU | 0.054 | 0.201 | 0.172 | 0.14 |
| TEDE-Sc2 | 0.04 | 0.154 | 0.162 | 0.136 |
| TEDE-aSPU2 | 0.047 | 0.191 | 0.148 | 0.131 |
| Scenario 3 (directional pleiotropy, InSIDE violated), $\beta = 0$ | | | | |
| LDA MR-Egger | 0.051 | 0.133 | 0.261 | 0.443 |
| PMR-Egger | 0.062 | 0.48 | 0.992 | 1 |
| TEDE-Sc | 0.038 | 0.738 | 0.995 | 1 |
| TEDE-aSPU | 0.04 | 0.806 | 0.998 | 1 |
| TEDE-Sc2 | 0.036 | 0.728 | 0.995 | 1 |
| TEDE-aSPU2 | 0.04 | 0.802 | 0.998 | 1 |
| Scenario 3 (directional pleiotropy, InSIDE violated), $\beta > 0$ ($h^2_{X\to Y} = 0.01$) | | | | |
| LDA MR-Egger | 0.065 | 0.107 | 0.227 | 0.389 |
| PMR-Egger | 0.057 | 0.456 | 0.99 | 1 |
| TEDE-Sc | 0.049 | 0.736 | 0.994 | 1 |
| TEDE-aSPU | 0.054 | 0.797 | 0.997 | 1 |
| TEDE-Sc2 | 0.04 | 0.688 | 0.991 | 1 |
| TEDE-aSPU2 | 0.047 | 0.783 | 0.998 | 1 |

**Table S3.** Rejection rates (type I error when there is 0 invalid IV; power otherwise) for testing valid IV assumptions. 1000 iterations. $p = 100$. High LD ($\rho = 0.7$).

| | 0 invalid | 10% invalid | 30% invalid | 50% invalid |
|---|---|---|---|---|
| Scenario 1 (balanced pleiotropy), $\beta = 0$ | | | | |
| LDA MR-Egger | 0.057 | 0.073 | 0.092 | 0.095 |
| PMR-Egger | 0.061 | 0.122 | 0.13 | 0.12 |
| TEDE-Sc | 0.04 | 0.196 | 0.194 | 0.183 |
| TEDE-aSPU | 0.052 | 0.28 | 0.248 | 0.255 |
| TEDE-Sc2 | 0.038 | 0.194 | 0.19 | 0.182 |
| TEDE-aSPU2 | 0.046 | 0.268 | 0.249 | 0.264 |
| Scenario 1 (balanced pleiotropy), $\beta > 0$ ($h^2_{\mathrm{X}\to\mathrm{Y}} = 0.01$) | | | | |
| LDA MR-Egger | 0.062 | 0.075 | 0.093 | 0.092 |
| PMR-Egger | 0.064 | 0.114 | 0.13 | 0.115 |
| TEDE-Sc | 0.049 | 0.201 | 0.2 | 0.185 |
| TEDE-aSPU | 0.056 | 0.245 | 0.231 | 0.239 |
| TEDE-Sc2 | 0.041 | 0.178 | 0.182 | 0.165 |
| TEDE-aSPU2 | 0.055 | 0.243 | 0.22 | 0.229 |
| Scenario 3 (directional pleiotropy, InSIDE violated), $\beta = 0$ | | | | |
| LDA MR-Egger | 0.057 | 0.222 | 0.495 | 0.607 |
| PMR-Egger | 0.061 | 0.803 | 1 | 1 |
| TEDE-Sc | 0.04 | 0.792 | 1 | 1 |
| TEDE-aSPU | 0.052 | 0.922 | 1 | 1 |
| TEDE-Sc2 | 0.038 | 0.782 | 1 | 1 |
| TEDE-aSPU2 | 0.046 | 0.92 | 1 | 1 |
| Scenario 3 (directional pleiotropy, InSIDE violated), $\beta > 0$ ($h^2_{\mathrm{X}\to\mathrm{Y}} = 0.01$) | | | | |
| LDA MR-Egger | 0.062 | 0.217 | 0.475 | 0.601 |
| PMR-Egger | 0.064 | 0.764 | 1 | 1 |
| TEDE-Sc | 0.049 | 0.783 | 1 | 1 |
| TEDE-aSPU | 0.056 | 0.911 | 1 | 1 |
| TEDE-Sc2 | 0.041 | 0.738 | 1 | 1 |
| TEDE-aSPU2 | 0.055 | 0.901 | 1 | 1 |

**Table S4.** Computation time of TEDE (seconds) averaged over 5 runs. TEDE-Sc gives the results for both TEDE-Sc and TEDE-Sc2, and TEDE-aSPU gives the results for both TEDE-aSPU and TEDE-aSPU2.

| | $p = 100$ | $p = 300$ | $p = 1000$ | $p = 2000$ |
|---|---|---|---|---|
| TEDE-Sc | 0.01 | 0.4 | 16.1 | 137 |
| TEDE-aSPU | 0.3 | 1.8 | 30.9 | 209 |

**Table S5.** Summary of the datasets used in the manuscript.

| Short Name | Reference and Link |
|---|---|
| ADNI | Shen et al. 2014<br>http://adni.loni.usc.edu/ |
| SCZ | Schizophrenia Working Group of the Psychiatric Genomics Consortium 2014<br>https://www.med.unc.edu/pgc/results-and-downloads |
| IGAP | Lambert et al. 2013<br>http://web.pasteur-lille.fr/en/recherche/u744/igap/igap_download.php |
| Lipid 2010 | Teslovich et al. 2010<br>http://csg.sph.umich.edu/abecasis/public/lipids2010 |
| Lipid 2013 | Willer et al. 2013<br>http://csg.sph.umich.edu/abecasis/public/lipids2013/ |
| FUSION | Gusev et al. 2016<br>http://gusevlab.org/projects/fusion/#reference-functional-data |

We also used the GWAS results based on imputed UK Biobank data (Sudlow et al. 2015; Neale Lab 2017) provided at http://biobank.ctsu.ox.ac.uk/crystal/.
